# Genetic Analysis of Methyl Anthranilate, Mesifurane, Linalool and Other Flavor Compounds in Cultivated Strawberry (*Fragaria* ×*ananassa*)

**DOI:** 10.1101/2020.10.07.330001

**Authors:** Christopher R. Barbey, Maxwell H. Hogshead, Benjamin Harrison, Anne E. Schwartz, Sujeet Verma, Youngjae Oh, Seonghee Lee, Kevin M. Folta, Vance M. Whitaker

**Affiliations:** Gulf Coast Research and Education Center, University of Florida, Wimauma, FL USA; Horticultural Sciences Department, University of Florida, Gainesville, FL USA

**Author notes:** Correspondence: Dr. Christopher Barbey, Horticultural Sciences Department, University of Florida, Gainesville FL 32611.

**Keywords:** Aromas, eQTL analysis, fruit volatiles, QTL analysis, terpenes, transcriptomics

## Abstract

The cultivated strawberry (*Fragaria* ×*ananassa*) is an economically important fruit crop that is intensively bred for improved sensory qualities. The diversity of fruit flavors and aromas in strawberry result mainly from the interactions of sugars, acids, and volatile organic compounds (VOCs) that are derived from diverse biochemical pathways influenced by the expression of many genes. This study integrates multi-omics analyses to identify QTL and candidate genes for multiple aroma compounds in a complex strawberry breeding population. Novel fruit volatile QTL were discovered for methyl anthranilate, methyl 2-hexenoate, methyl 2-methylbutyrate, mesifurane, and a shared QTL on Chr 3 was found for nine monoterpene and sesquiterpene compounds, including linalool, 3-carene, β-phellandrene, α-limonene, linalool oxide, nerolidol, α-caryophellene, α-farnesene, and β-farnesene. Fruit transcriptomes from a subset of sixty-four individuals were used to support candidate gene identification. For methyl esters including the grape-like methyl anthranilate, a novel *ANTHANILIC ACID METHYL TRANSFERASE*–like gene was identified. Two mesifurane QTL correspond with the known biosynthesis gene *O-METHYL TRANSFERASE 1* and a novel *FURANEOL GLUCOSYLTRANSFERASE*. The shared terpene QTL contains multiple fruit-expressed terpenoid pathway-related genes including *NEROLIDOL SYNTHASE 1* (*FanNES1*). The abundance of linalool and other monoterpenes is partially governed by a co-segregating expression-QTL (eQTL) for *FanNES1* transcript variation, and there is additional evidence for quantitative effects from other terpenoid-pathway genes in this narrow genomic region. These QTL present new opportunities in breeding for improved flavor in commercial strawberry.

## INTRODUCTION

The dessert strawberry (*Fragari*a × *ananassa*) is a widely celebrated fruit with increasing consumption. For decades consumers have reported the desire for improved flavor in commercial strawberry (Fletcher 1917; Chambers 2013). The aroma intensity of modern cultivars is lower than in wild strawberries (Ulrich and Olbricht 2014), and breeding efforts seek to reclaim these qualities. Today flavor and aroma are central priorities of strawberry breeding programs (Faedi et al. 2002; Whitaker et al. 2011; Vandendriessche et al. 2013). However, breeders face a significant challenge in the recapture and consolidation of genetics contributing to favorable flavors and aromas. Genetic and genomic analysis has been used to identify these elements and contribute to the breeding of new cultivars with improved sensory qualities.

Strawberry flavor and aroma are dictated by several factors, including sugars and acids. But it is the trace volatile organic compounds (VOCs) that shape the sensory experience (Bood and Zabetakis 2002). Volatile organic compounds, represented broadly as esters, alcohols, terpenoids, furans and lactones, are a substantial portion of the fruit secondary metabolome and contribute to aroma, flavor, disease resistance, pest resistance and overall fruit quality (Ulrich et al. 1997; Arroyo et al. 2007). Various studies have helped to identify human preferences for individual strawberry aroma and flavor compounds (Schwieterman et al. 2014; Ulrich et al. 1997; Schieberle and Hofmann 1997; Larsen and Poll 1992). Of hundreds of strawberry VOCs, these studies agree on fewer than ten that clearly influence human preference (Schwieterman et al. 2014). Introgressing important compounds into commercially viable cultivars has been aided by efforts in volatilomics QTL detection (Urrutia et al. 2017), multi-omics identification of VOC candidate genes (Pillet et al. 2017; Chambers et al. 2014), integration of sensory and consumer preference data (Schwieterman et al. 2014; Sánchez-Sevilla et al. 2014), and ultimately introgression of genes via marker-assisted selection (Folta and Klee 2016; Rambla et al. 2017; Eggink et al. 2014).

Only a few strawberry genes controlling desirable aroma compounds have been identified with confidence. These include biosynthesis genes for linalool (Aharoni et al. 2004), mesifurane (Wein et al. 2002), γ-decalactone (Chambers et al. 2014; Sánchez-Sevilla et al. 2014) and methyl anthranilate (Pillet et al. 2017). The gene *ANTHRANILIC ACID METHYL TRANSFERASE* (*FanAAMT*), located on octoploid chromosome group 4, is a necessary-but-not-sufficient gene for catalyzing the methylation of anthranilate into the grape-like aroma compound methyl anthranilate (Pillet et al. 2017). Methyl anthranilate production has been long regarded as a complex trait, governed by multiple genes and strong environmental influences. Methyl anthranilate is produced abundantly in the fruit of the diploid strawberry sp. *Fragaria vesca*, but it is reported in only a few octoploid varieties including ‘Mara des Bois’ and ‘Mieze Schindler’ (Ulrich and Olbricht 2016). For terpenoid biosynthesis, strawberry *NEROLIDOL SYNTHASE 1 (NES1)* was identified by comparing diploid and octoploid species, which are enriched respectively for nerolidol or linalool. A truncated plastid targeted signal in the octoploid *FanNES1* gene retargets the enzyme to the cytosol, where there is abundant precursor for linalool biosynthesis (Aharoni et al. 2004). Recent reports have complicated this story somewhat, as three octoploid *F. virginiana* lines, hexaploid *F. moschata*, and some diploid strawberries produce linalool without this truncation (Ulrich and Olbricht 2013). In mesifurane biosynthesis, the gene *O-METHYL TRANSFERASE 1* (*FanOMT1*) catalyzes the methylation of furaneol to create mesifurane (Wein et al., 2002, Zorrilla-Fontanesi et al., 2012). Mesifurane abundance is affected by a common *FanOMT1* promoter loss-of-function allele, which both eliminates gene expression and mesifurane production. Only one copy of the competent *FanOMT1* allele is reportedly sufficient for robust production, however a lack of production is sometimes observed even in the homozygous positive state (Cruz-Rus et al. 2017). An octoploid gene encoding *QUINONE REDUCTASE* (*FanQR*) can produce furaneol *in vitro*, however no natural variants of this gene have been established which vary mesifurane levels *in vivo* (Raab et al. 2006). Similarly, glucosylation of both furaneol and mesifurane are known to occur in strawberry, however genetic variation has not been established for this step. Several furaneol glucosyltransferases have been cloned and characterized *in vitro* from *F. ×ananassa* (Song et al. 2016; Yamada et al. 2019).

This research integrates high-density genotyping and non-targeted fruit volatile metabolomics from eight pedigree-connected octoploid crosses (n=213) (Figure S1). Fruit transcriptomes from a subset of individuals (n=61) were used to identify fruit-expressed candidate genes within QTL regions (Figure S1). Three maps were utilized independently in this analysis, as less than one-third of octoploid subgenome-specific markers are incorporated in any single octoploid genetic map (Anciro et al. 2018; van Dijk et al. 2014). These are the ‘Holiday’ × ‘Korona’ (van Dijk et al. 2014) and FL_08-10 × 12.115-10 (Verma et al. 2017) genetic maps, and the *F. vesca* physical map. The correspondence of ‘Holiday’ × ‘Korona’ linkage groups to the recent octoploid ‘Camarosa’ reference genome (Edger et al. 2019) helped specify the subgenomic identity of QTL (Hardigan et al. 2020). To correspond QTL markers to specific candidate gene regions, marker nucleotide sequences from the IStraw35 SNP genotyping platform were aligned by sequence to the octoploid genome. However, the very high sequence identity between homoeologous chromosomes limited the specificity of this approach. Evidence from all of these resources were integrated to specify candidate gene regions.

Fruit transcriptomes were used to identify expressed genes within QTL regions and to associate trait/transcript levels. Genotypic data were associated with fruit transcriptomics data via expression-QTL (eQTL) analysis. Transcript eQTL analysis identifies genetic variants associated with heritable transcript level variation. Transcript eQTL often correspond to the locus of the originating gene (*cis*-eQTL), and often signify gene promoter mutation or gene presence/absence variation. In cases where a trait is governed by simple genetic control of transcript levels of a causal gene, an eQTL should be detected which co-segregates with trait QTL markers. This approach can help specify the casual mechanisms behind trait QTL. Previous eQTL analyses in these same fruit RNA-seq populations identified hundreds of fruit eQTL, the vast majority of which were proximal to the originating gene locus (Barbey et al. 2020). Global transcriptomes from tissues throughout the octoploid cultivar ‘Camarosa’ were used to correlate candidate genes and transcript abundance with ripening-associated volatile biosynthesis (Sánchez-Sevilla et al. 2017).

## MATERIALS AND METHODS

### Plant Materials

Eight controlled crosses were made among octoploid cultivars and also elite breeding lines from the University of Florida. These were ‘Florida Elyana’ × ‘Mara de Bois’ (population 10.113), ‘Mara des Bois’ × ‘Florida Radiance’ (population 13.75), ‘Strawberry Festival’ × ‘Winter Dawn’ (population 13.76), 12.115-10 × 12.121-5 (population 15.89), 12.22-10 × 12.115-10 (population 15.91), 12.115-10 × 12.74-39 (population 15.93), ‘Florida Elyana’ × 12.115-10 (population 16.11), and ‘Mara des Bois’ × ‘Mara des Bois’ (population 16.85). Seedlings were clonally propagated by runners in a summer nursery to generate multiple plants (clonal replicates), and two to four plants representing original seedlings were established in single plots in the fruiting field. (Barbey et al. 2020).

### Field Collection

All fruit were harvested from a field maintained under commercial growing practices during winter growing seasons at the Gulf Coast Research and Education Center (GCREC) in Wimauma, Florida. Fruit from populations 15.89, 15.91, 15.93 were harvested on February 19, March 15, and April 14, 2016. Populations 16.11, 16.85, were harvested on February 16 and March 2, 2017. Populations 13.75 and 13.76 were harvested during the winter of 2014. Population 10.133 was sampled on January 20, February 11, February 25, and March 18, 2011 (Chambers 2013). Harvest days were selected based on dry weather and moderate temperature, both on the day of harvest and for several days preceding harvest to maximize volatile production and capture. Three mature fruit per genotype were cleaned, collected into a single sample bag, crushed, and immediately flash-frozen in liquid nitrogen, in the field. Samples were transported to Gainesville, Florida and maintained at -80 °C.

### Sample Processing and Preparation

Unprocessed frozen fruit samples were equilibrated from -80 °C to liquid nitrogen temperature before being pureed in an electric blender. Frozen puree was collected into a 50ml sterile collection tube and stored at -80 °C. For volatile sample processing, 3g of fruit puree from each sampled genotype at a single timepoint was aliquoted into two technical replicate 20ml headspace vials and combined with 3ml 35% NaCl solution containing 1ppm 3-hexanone as an internal standard. Prepared vials were stored at -80°C, thawed at room temperature and vortexed prior to GC-MS analysis.

### Volatile Metabolomic Profiling and Analysis

Samples were equilibrated to 40 °C for 30 min in a 40 °C heated chamber. A 2 cm tri-phase SPME fiber (50/30 μm DVB/Carboxen/PDMS, Supelco, Bellefonte, PA, USA) was exposed to the headspace for 30 min at 40°C for volatile collection and concentration. The fiber was then injected into an Agilent 6890 GC (for 5 min at 250°C for desorption of volatiles. Inlet temperatures were maintained at 250°C, ionizing sources at 230°C, and transfer line temperatures at 280°C. The separation was performed via DB5ms capillary column (60 m x 250 μm x 1.00 μm) (J&W, distributed by Agilent Technologies) at a constant flow (He: 1.5 ml per min). The initial oven temperature was maintained at 40°C for 30 seconds, followed by a 4°C per min increase to a final temperature of 230°C, then to 260°C at 100°C per min, with a final hold time of 10 min. Data were collected using the Chemstation G1701 AA software (Hewlett-Packard, Palo Alto, CA).

Chromatograms were processed using the Metalign metabolomics preprocessing software package (Lommen and Kools 2012). Baseline and noise corrections were performed using a peak slope factor of 1x noise, and a peak threshold factor of 2x noise. Auto-scaling and iterative pre-alignment options were not selected. A maximum shift of 100 scans before peak identification, and 200 scans after peak identification was used. In later validation steps, these search tolerances were determined to be sufficiently inclusive while also limiting to false-positives. The MSClust software package was then used for statistical clustering of ions based on retention time and co-variance across the population using default parameters (Tikunov et al. 2012). Clusters were batch queried against the NIST08 reference database using Chemstation G1701 AA software (Hewlett-Packard, Palo Alto, CA). Library search outputs were parsed using a custom Perl script prior to multivariate analysis. Chromatograms were batched by GC-MS sampling year to mitigate ion misalignments caused by system-dependent retention time shifts. VOC abundances were normalized based on the internal standard. Normalized data between seasons were consolidated manually based on elution order, NIST identification, and rerunning of sample standards.

### Genotyping of Flavor and Aroma Populations

Individuals from all populations were genotyped using the IStraw90 (Bassil et al. 2015)) platform, excepting populations 16.11 and 16.89 which were genotyped using the IStraw35 platform (Verma et al. 2017). All parents and 204 progeny were selected for genotyping based on the segregation of desirable fruit volatiles. Sequence variants belonging to the Poly High Resolution (PHR) and No Minor Homozygote (NMH) marker classes were included for association mapping. Mono High Resolution (MHR), Off-Target Variant (OTV), Call Rate Below Threshold (CRBT), and other marker quality classes, were discarded and not used for mapping. Individual marker calls inconsistent with Mendelian inheritance from parental lines were removed.

### Fruit Transcriptome Assembly and Analysis

Mature fruit from 61 parents and progeny from the biparental populations 10.113, 13.75, and 13.76 were sequenced via Illumina paired-end RNA-seq (avg. 65million 2×100bp reads) as previously described (Barbey et al. 2019). Briefly, RNA-seq reads were assembled based on the *Fragaria* ×*ananassa* octoploid ‘Camarosa’ annotated genome, with reads mapping equally-well to multiple loci discarded from the analysis. Separately, raw RNA-seq reads from the ‘Camarosa’ strawberry gene expression atlas study (Sánchez-Sevilla et al. 2017) were assembled via the same method. Transcript abundances were calculated in Transcripts Per Million (TPM). Fruit eQTL analysis was performed using the mixed linear model method implemented in GAPIT v3 (Tang et al. 2016) as described in (Barbey et al. 2020).

### Genetic Association of Fruit Volatiles

Relative volatile abundance values were rescaled using the Box-Cox transformation algorithm (Box and Cox 1964) performed in R (R. Development Core Team 2014)using R-studio (Racine 2011) prior to genetic analysis. GWAS on fruit volatiles was performed using the mixed linear model method implemented in GAPIT v3 (Tang et al. 2016) in R, using marker positions oriented to the *F. vesca* diploid physical map. Significantly-associated volatiles were then reanalyzed in GAPIT using the ‘Holiday’ × ‘Korona’ and FL_08-10 × 12.115-10 genetic maps. Metabolomic associations were evaluated for significance based on the presence of multiple co-locating markers of *p*-value < 0.05 after FDR multiple comparisons correction (Benjamini and Hochberg 1995). Narrow-sense heritability (*h*^*2*^) estimates were derived from GAPIT v3 while single-marker analysis was performed via ANOVA in R to investigate allelic effects.

### Analysis of Candidate Genes

All gene models in the ‘Camarosa’ genome were analyzed with the BLAST2GO pipeline and the Pfam protein domain database. Genes with significant homology to known volatile biosynthesis genes including *FanOMT* and *FanAAMT* were collected from the ‘Camarosa’ genome using BLAST with inclusive criteria. This process was replicated for candidate genes including anthranilate synthase alpha subunit (*FanAS-α)* and others not presented in this analysis. Deduced protein sequences from transcripts were aligned using the slow progressive alignment algorithm in the CLC Genomics Workbench 11 (Gap Open cost=10; Gap Extension=1). Tree construction was performed using the Neighbor Joining method with Jukes-Cantor distance measuring with 1,000 bootstrapping replicates. Fruit transcript heatmaps were added to the cladogram to show the maximum transcript level detected among the 61 fruit transcriptomes. Genes putatively belonging to published volatile biosynthesis gene families were selected for fruit transcript eQTL analysis, using methods described previously (Barbey et al. 2020; Barbey et al. 2019). The 200 genes surrounding the most-correlated volatile QTL markers were also analyzed for eQTL and compared for co-segregation with volatile QTL.

### High Resolution Melting Marker Test

For the marker test of mesifurane, two validation crosses were created consisting of ‘Florida Beauty’× 15.89-25 (population 18.50, n= 27) and 15.34-82 × 15.89-25 (population 18.51, n=44). Total genomic DNA was extracted using the simplified cetyltrimethylammonium bromide (CTAB) method described by (Noh et al. 2018) with minor modifications. To develop HRM markers for mesifurane, two probes were selected (AX-166520175, AX-166502845) that were highly associated with mesifurane on Chr 1 QTL. The primers 5’-CCCTTGGCATCAATATTTGTGAAT-3’ and 5’-GAACTCCATTAGAAATCAAGTTATCAGC-3’ were designed for AX-166520175, and the 5’-CTGATCCTGCTTCAAGTACAAG-3’ and 5’-TCAATGAAGACACTTGATCGAC-3’ were designed for AX-166502845 using IDT’s PrimerQuest Software (San Jose, CA, USA). PCR amplifications were performed in a 5 µl reaction containing 2× AccuStart^™^ II PCR ToughMix® (Quantabio, MA, USA), 1× LC Green® Plus melting dye (BioFire, UT, USA), 0.5 µM of each HRM primer sets and 1 µl of DNA. The PCR and HRM analysis were performed in a LightCycler® 480 system II (Roche Life Science, Germany) using a program consisting of an initial denaturation at 95°C for 5 minutes; 45 cycle of denaturation at 95°C for 10 seconds, annealing at 62°C for 10 seconds, and extension at 72°C for 20 seconds. After PCR amplification, the samples were heated to 95°C for 1 min and cooled to 40°C for 1 min. Melting curves were obtained by melting over the desired range (60 - 95°C) at a rate of 50 acquisitions per 1°C. Melting data were analyzed using the Melt Curve Genotyping and Gene Scanning Software (Roche Life Science, Germany). Analysis of HRM variants was based on differences in the shape of the melting curves and in *Tm* values.

## RESULTS

### Strawberry Fruit Flavor and Aroma QTL

Volatile aroma QTL were discovered for methyl anthranilate (CAS 134-20-3), methyl 2-methylbutyrate (868-57-5), methyl 2-hexenoate (CAS 2396-77-2) (Figures 1 and 2), mesifurane (CAS 4077-47-8) (Figure 3), and nine mono- and sesquiterpenes compounds (Figures 4 and 5). These terpenes include linalool (CAS 78-70-6), 3-carene (CAS 498-15-7), β-phellandrene (CAS 555-10-2), α-limonene (CAS 5989-27-5), linalool oxide (CAS 60047-17-8), nerolidol (CAS 7212-44-4), α-caryophellene (CAS 6753-98-6), α-farnesene (CAS 502-61-4), and β-farnesene (CAS 77129-48-7). Many of these compounds are consensus determinants of preferred strawberry flavor in human sensory trials, including grape-like methyl anthranilate (Ulrich et al. 1997), fruity-sweet methyl 2-methylbutanoate (Schieberle and Hofmann 1997), fruity linalool (Larsen and Poll 1992; Schieberle and Hofmann 1997; Ulrich et al. 1997; Schwieterman 2013), and sherry/caramel-like mesifurane (Larsen and Poll 1992; Schieberle and Hofmann 1997; Ulrich et al. 1997; Schwieterman 2013). Population-wide fruit transcript-level data for all candidate genes in the following analysis are provided in Table S1, and transcript levels throughout the ‘Camarosa’ plant are provided in Table S2.

**Figure 1.**
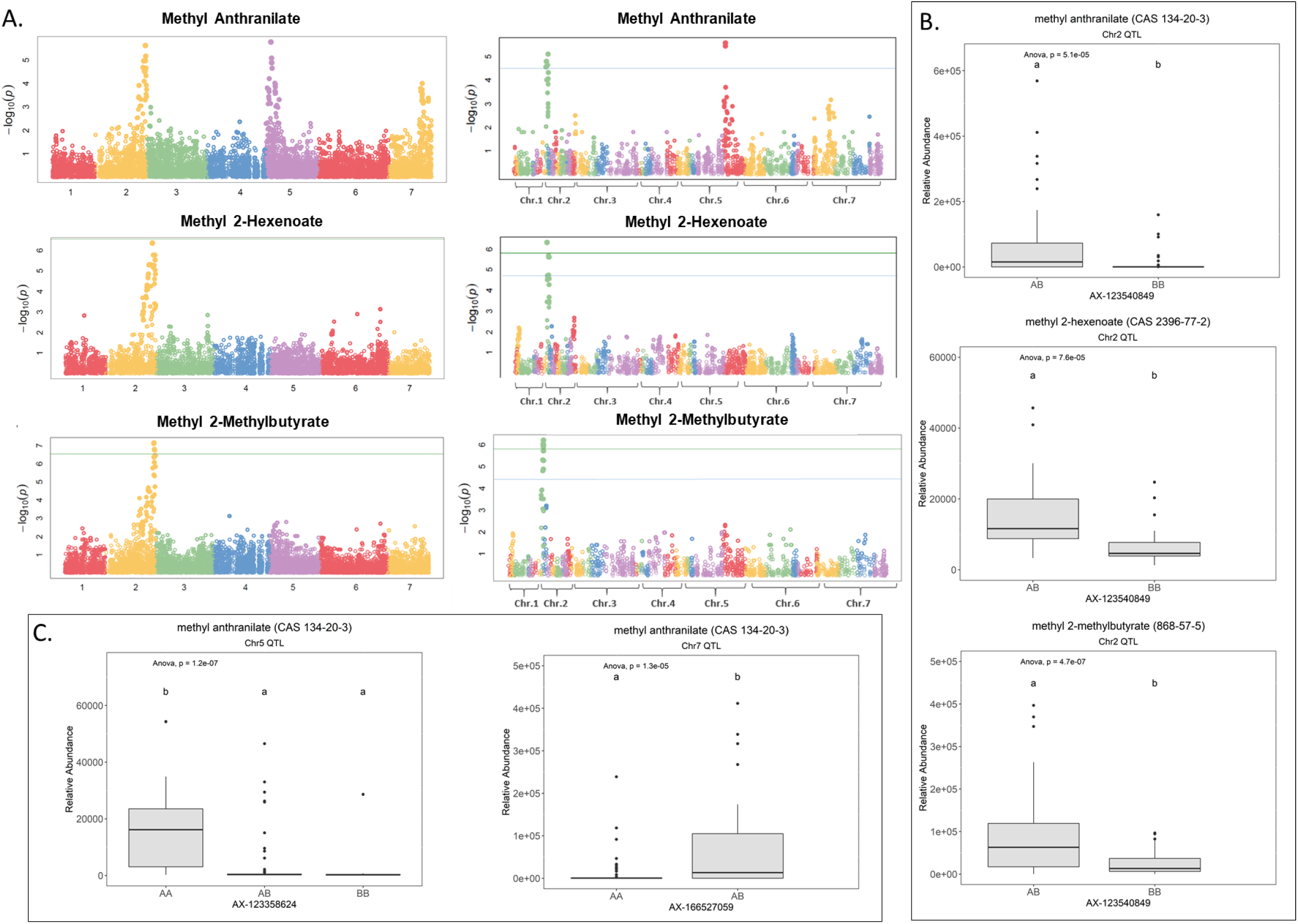
Methyl Ester QTL in Strawberry Fruit. **(A)** QTL Manhattan plots for Methyl Anthranilate, Methyl 2-hexenoate, and Methyl 2-Methylbutryrate using the *F. vesca* physical map (left column) and the FL_08-10 × 12.115-10 genetic map (right column). **(B)** Effect-sizes of a shared methyl ester QTL marker on Chr 2. Lower-case letters indicate statistically significance mean differences at *p* < 0.05 (ANOVA). **(C)** Effect-sizes for the methyl anthranilate QTL on Chr 5 and the putative Chr 7 QTL.

**Figure 2.**
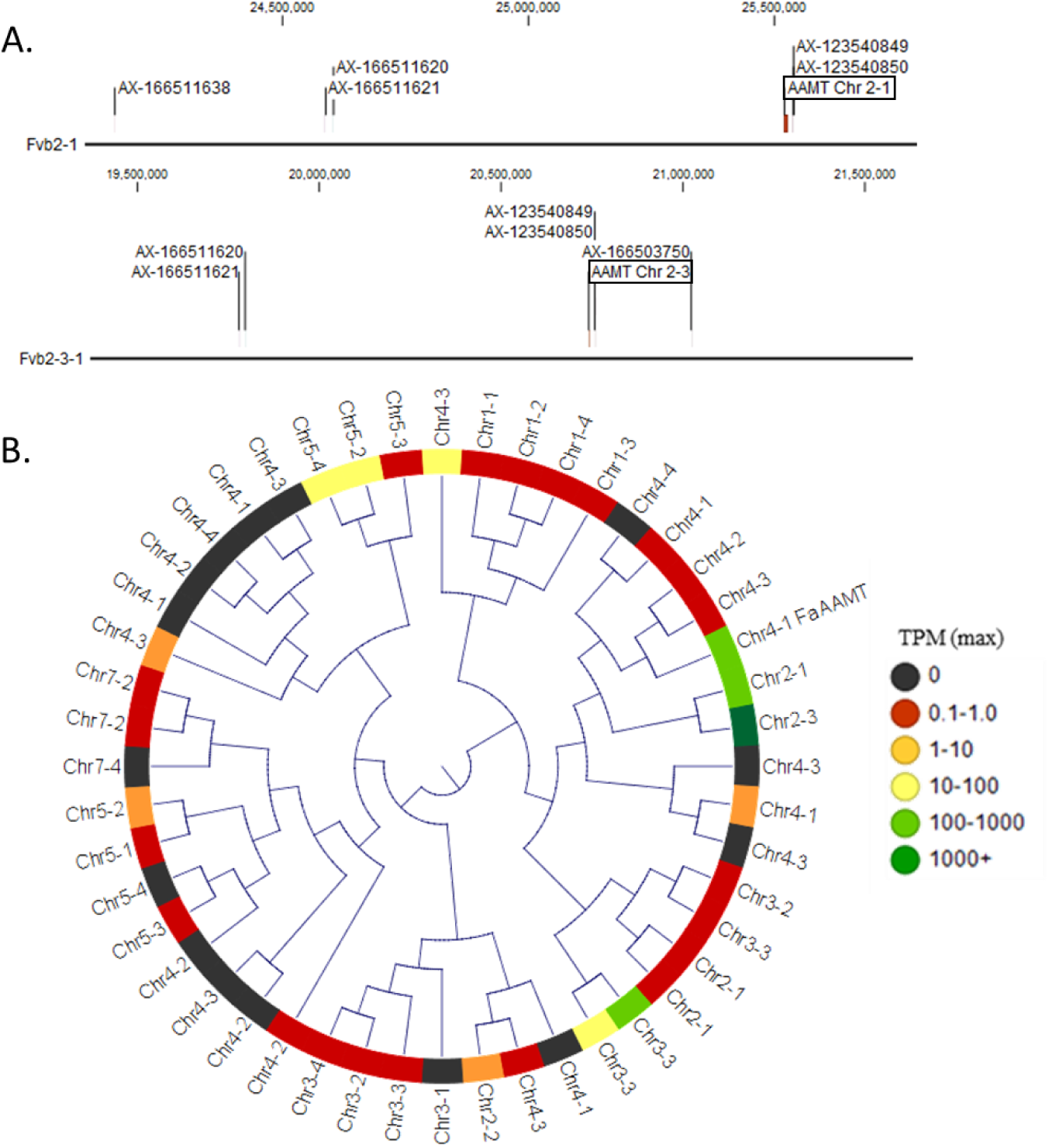
Methyl Ester and Methyl Anthranilate Candidate Genes. **(A)** The Chr 2 Methyl Ester QTL markers correspond to two homoeologous physical regions containing Anthranilic Acid Methyl Transferase-like (*AAMT-like*) genes on Chr 2-1 (top) and Chr 2-3 (bottom). **(B)** *AAMT-like* deduced proteins in the ‘Camarosa’ genome are shown in a neighbor-joining cladogram, with transcript abundance heatmaps representing the highest TPM detected among the fruit transcriptomes. The Chr 2-1 and Chr 2-3 AAMT-like candidates are highly identical to the published AAMT-like ‘Camarosa’ homolog (Chr 4-1 *FanAAMT*), and are highly abundant transcripts in the fruit.

**Figure 3.**
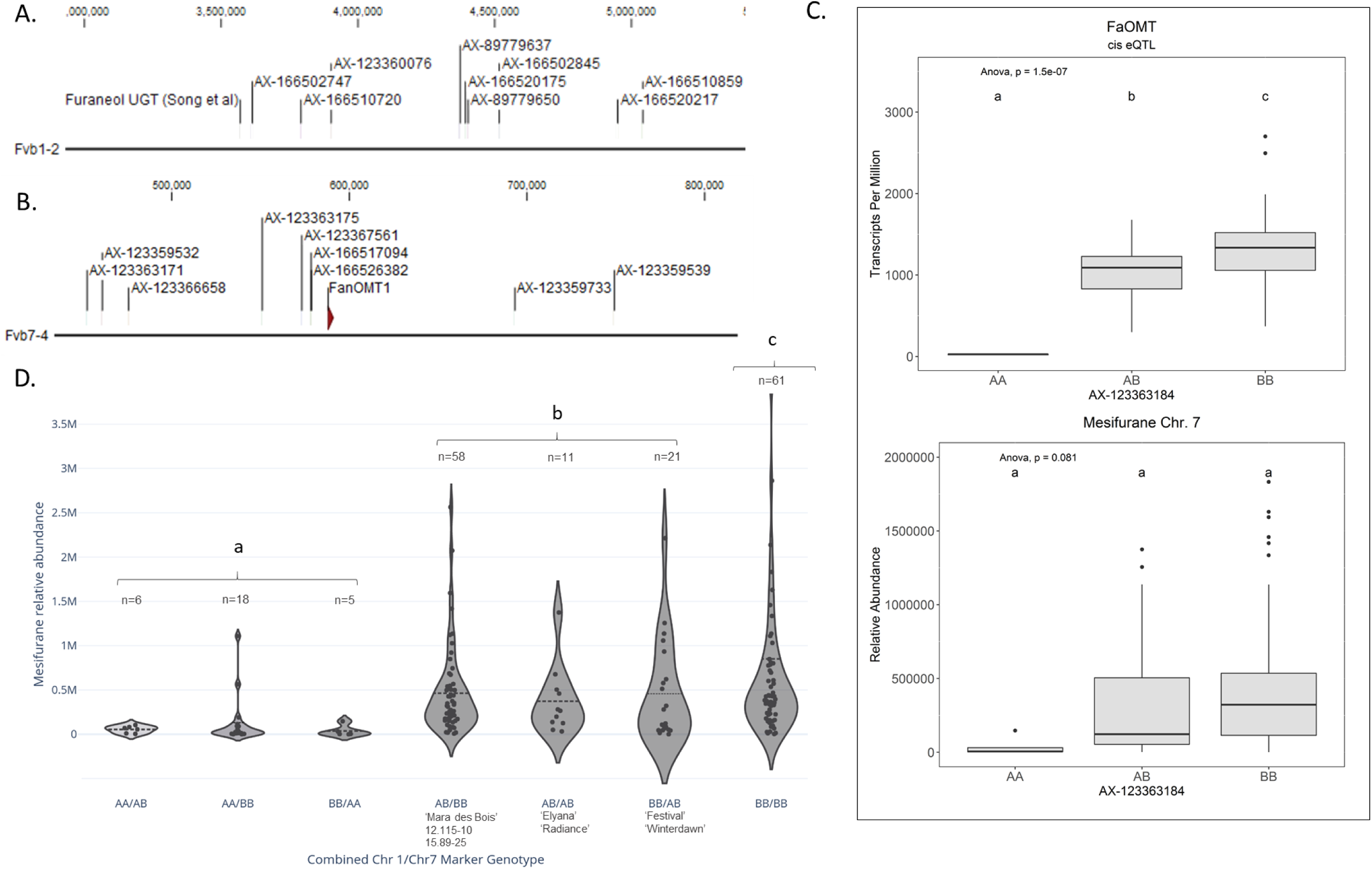
Mesifurane QTL and Candidate Genes. **(A)** The Chr 1 Mesifurane QTL markers correspond to a region containing a characterized Furaneol UGT homolog. **(B)** The Chr 7 Mesifurane QTL markers correspond to a region containing the published mesifurane *FanOMT1* biosynthesis gene. **(C)** A cis-eQTL was detected for *FanOMT1* (top), a gene known to be transcriptionally variable due to a common allele. The *FanOMT1* eQTL co-segregates with mesifurane (bottom) but is confounded by additional factors. Lower-case letters indicate statistically significance mean differences at *p* < 0.05 (ANOVA). **(D)** Allelic combinations at the Chr 1 and Chr 7 markers demonstrate an epistatic effect between the two loci. At least one competent allele at each loci is required for robust production of mesifurane (AA/BB vs. AB/BB Tukey HSD *p* = 2.4e-7), while the double-homozygous allelic state produces statistically elevated mesifurane levels (BB/BB vs. AB/AB, AB/BB, and BB/AB ANOVA (F(1,149) = 4.38, *p* = 0.038). The Chr 1/Chr 7 genotypes in parental lines is shown with the *n* of each allelic category. Dotted lines represent the population means. Lower-case letters indicate statistically significance mean differences at *p* < 0.05 (ANOVA).

**Figure 4.**
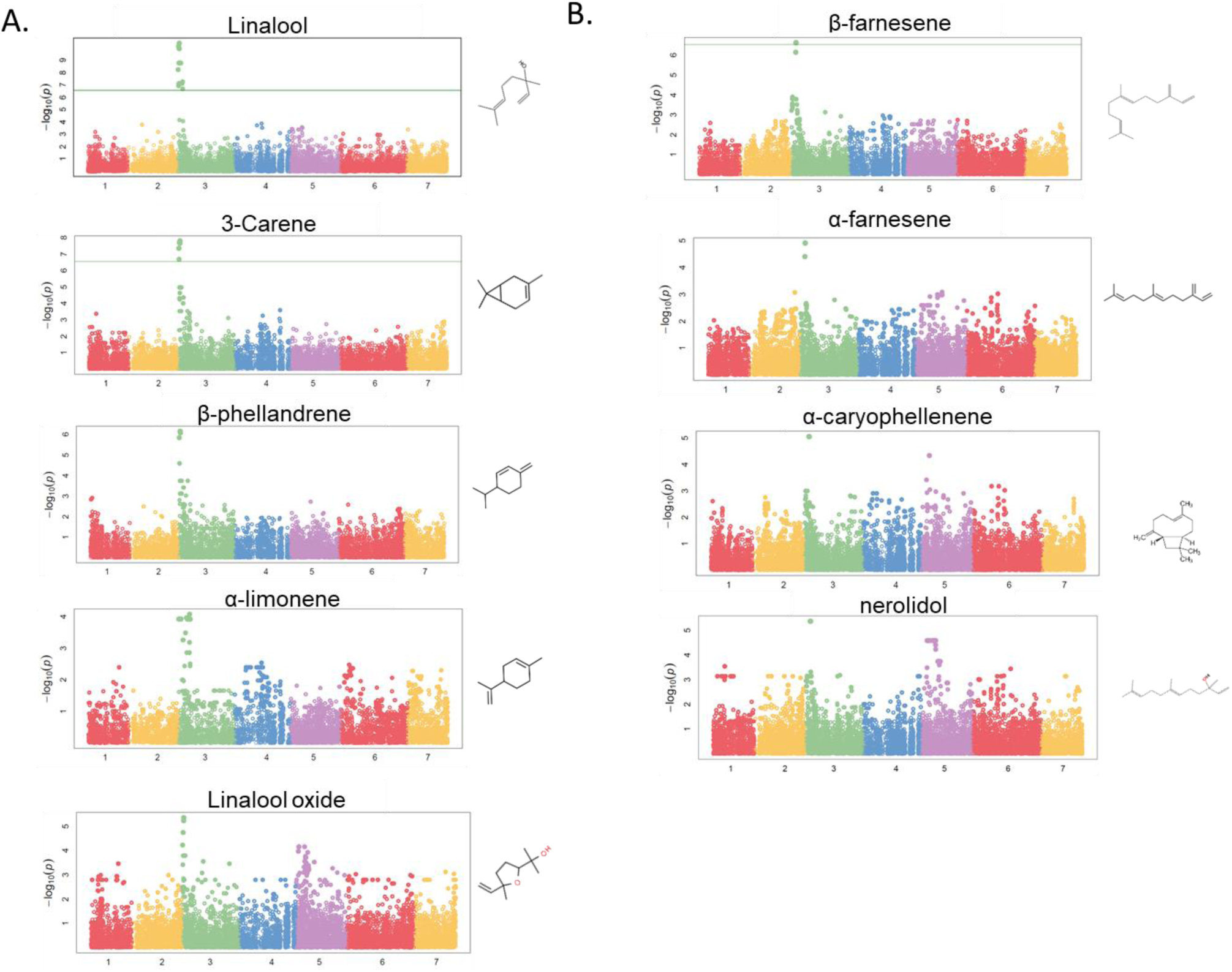
Terpene Volatile QTL in Strawberry Fruit. **(A)** QTL Manhattan plots for the monoterpenes linalool (*r*^*2*^ = 0.15, *p* = 4.2e-11), 3-carene (*r*^*2*^ = 0.09, *p* = 1.2e-8), β-phellandrene (*r*^*2*^ = 0.21, *p* = 7.0e-7), α-limonene (*r*^*2*^ = 0.06, *p* = 7.0e-7), linalool oxide (*r*^*2*^ = 0.08, *p* = 4.4e-6). **(B)** the sesquiterpenes β-farnesene (*r*^*2*^ = 0.12, *p* = 2.4e-7), α-farnesene (*r*^*2*^ = 0.16, *p* = 1.6e-6), α-caryophellenene (*r*^*2*^ = 0.10, *p* =8.9e-6), and nerolidol (*r*^*2*^ = 0.09, *p* = 4.3e-6). Chemical structures are shown for each compound.

**Figure 5.**
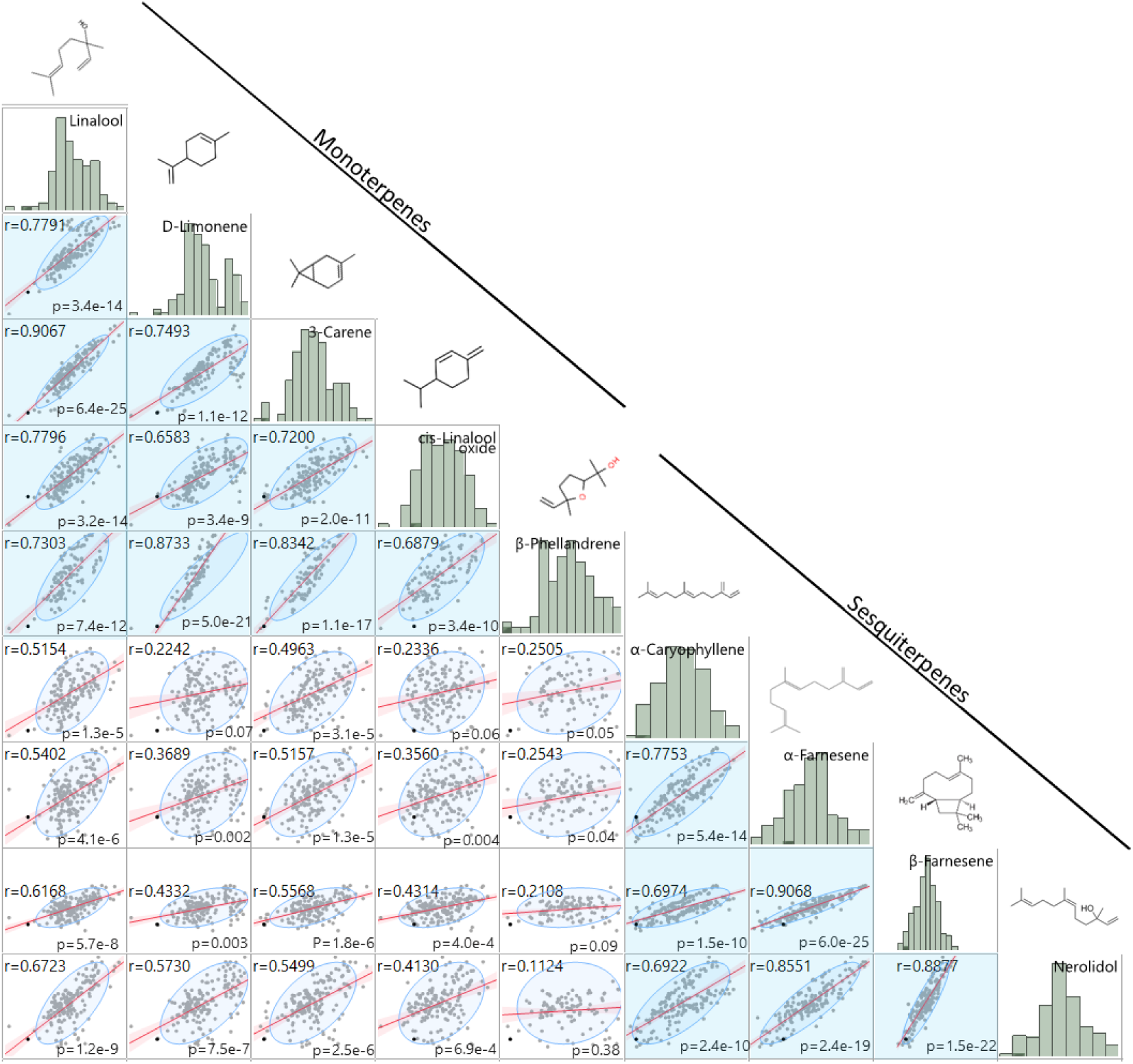
Comparison of Terpene Fruit Volatile Abundances. Mono- and sesquiterpene volatile abundances are strongly positively correlated within-group (blue shade) and less strongly positively correlated between groups (white shade). Regression statistics are shown for each pairwise comparison. Histograms of log2-transformed GC/MS relative-abundance values and chemical structures are denoted above each compound.

### Methyl Anthranilate and Methyl Ester QTL and Candidate Genes

Methyl anthranilate (*h*^*2*^= 0.59) QTL were identified on octoploid linkage groups (LGs) 2A and 5A of the FL_08-10 × 12.115-10 map (Figure 1A, Table 1). The LG 2A QTL is shared with methyl 2-hexenoate (*h*^*2*^ =0.43) and methyl 2-methylbutyrate (*h*^*2*^ =0.79) (Figure 1A; Table 1). This shared methyl ester QTL accounts for 11.7% of methyl anthranilate variance (*p*= 5.1e-5), 22.8% of methyl 2-hexenoate variance (*p*= 7.6e-5), and 18.1% of methyl 2-methylbutyrate variance *(p*= 4.7e-7) (AX-123540849, single-marker analysis) (Figure 1B). The QTL on LG 5D explains 19.7% of methyl anthranilate variance (AX-123358624, *p*= 6.324e-07). A diagram of known and hypothesized methyl anthranilate pathway components is provided in Figure S2.

**Table 1.** Methyl Anthranilate QTL and Candidate Gene Positions.

| Chr 2 QTL | *Methyl Anthranilate* *p*-value | methyl 2-hexenoate *p*-value | methyl 2-methylbutyrate *p*-value | Holiday x Korona map |  | 14.95 map |  | 'Camarosa' genome physical position |  |  |  |
| --- | --- | --- | --- | --- | --- | --- | --- | --- | --- | --- | --- |
|  |  |  |  | LG | Position | LG | Positon | Chr 2-1 | Chr 2-2 | Chr 2-3 | Chr 2-4 |
| *AAMT 2-1 candidate* | - | - | - | - | - | - | - | 25,519,186 | - | - | - |
| *AAMT 2-3 candidate* | - | - | - | - | - | - | - | - | - | 3,328,115 | - |
| AX-123540849 | 2.46E-06 | 1.72E-06 | 7.08E-08 | - | - | 2A | 9.799 | 25,537,654 | - | 3,313,464 | 25,836,027 |
| AX-166503750 | 6.82E-06 | 1.72E-06 | 8.49E-07 | - | - | 2A | 7.05 | - | - | 3,048,471 | 26,031,849 |
| AX-123540850 | 1.07E-05 | 2.47E-05 | 2.99E-04 | - | - | 2A | 10.409 | 25,537,654 | - | 3,313,464 | 25,836,027 |
| AX-123360428 | 1.09E-05 | 6.36E-06 | 6.74E-05 | - | - | - | - | - | - | - | 25,369,790 |
| AX-166511620 | 1.09E-05 | 6.36E-06 | 6.74E-05 | 2A | 78.514 | 2A | 1.851 | 24,602,373 | - | 4,275,047 | 25,558,703 |
| AX-166511621 | 1.09E-05 | 6.36E-06 | 6.74E-05 | - | - | - | - | 24,586,556 | - | 4,292,083 | 25,573,913 |
| AX-166503618 | 2.05E-05 | 4.50E-07 | 2.23E-04 | - | - | 2A | 0.304 | - | - | - | - |
| AX-166511638 | 2.05E-05 | 4.50E-07 | 2.23E-04 | - | - | - | - | 24,157,920 | - | - | - |
| AX-166511640 | 2.05E-05 | 4.50E-07 | 2.23E-04 | - | - | 2A | 0.304 | - | - | - | - |
| AX-166511795 | 2.73E-05 | 2.95E-06 | 4.88E-05 | - | - | 2A | 9.647 | - | - | - | 26,103,640 |
| Chr 5 QTL | Methyl Anthranilate *p*-value |  |  | LG | Position | LG | Positon | Chr 5-1 | Chr 5-2 | Chr 5-3 | Chr 5-4 |
| *GTP 5-4 candidate* | - |  |  | - | - | - | - | - | - | - | 1,182,596 |
| AX-123358608 | 3.63E-06 |  |  | 5D | 6.757 | - | - | 2,613,648 | - | 24,857,145 | 2,876,834 |
| AX-166506768 | 3.63E-06 |  |  | 5D | 6.757 | - | - | 2,524,864 | 3,118,449 | - | - |
| AX-166524167 | 3.63E-06 |  |  | 5D | 6.757 | 5D | 12.114 | 2,568,242 | 3,075,907 | - | - |
| AX-123361979 | 5.35E-06 |  |  | 5D | 8.109 | 5D | 12.114 | 2,212,147 | 3,426,121 | 24,503,708 |  |
| AX-166514937 | 9.60E-05 |  |  | 5D | 9.484 | - | - | 3,950,769 | - | 24,414,731 | 3,354,090 |
| AX-89892771 | 9.60E-05 |  |  | 5D | 9.484 | - | - | 3,984,739 |  | 24,382,881 | 3,363,253 |
| AX-166506799 | 1.51E-04 |  |  | 5D | 10.802 | - | - | - | - | 22,754,788 | 4,769,438 |
| AX-166518037 | 2.97E-04 |  |  | 5D | 1.351 | 5D | 8.12 | - | - | - | 1,615,784 |
| AX-166505352 | 3.28E-04 |  |  | 5D | 9.484 | - | - | 4,160,836 | - | - |  |
| AX-123358650 | 1.59E-03 |  |  | 5D | 9.484 | - | - | 5,695,643 | 4,386,116 | - | - |

The QTL on LG 2A broadly corresponds to the *F. vesca*-like Chr 2-2 of the ‘Camarosa’ octoploid genome, based on chromosome-wide genetic-genomic connections (Hardigan et al. 2020). However, nucleotide BLAST of IStraw35 methyl anthranilate marker sequences align non-specifically to all ‘Camarosa’ Chr 2 homoeologs except Chr 2-2 (Table 1). The corresponding syntenic regions of Chr 2-1 and Chr 2-3 each contain an *ANTHRANILIC ACID METHYL TRANSFERASE*-like homoeolog. The most-correlated QTL marker (AX-123540849) aligns 10 kb (2 genes) from the *AAMT-like* gene on Chr 2-1 (maker-Fvb2-1-snap-gene-255.58), and 14.6 kb (4 genes) from the *AAMT-like* gene on Chr 2-3 (maker-Fvb2-3-snap-gene-33.59) (Figure 2A and B; Table 1). These two *FanAAMT*-*like* genes have the highest sequence identity to the published Chr 4 *FanAAMT* gene excepting the highly-expressed *FanAAMT* gene on Chr 4-1 and its non-expressed homoeologs (Figure 2C). Reference-based RNA-seq shows both candidate *FanAAMT*-like transcripts are highly abundant in some fruit transcriptomes, but relatively low or almost absent in others (Chr 2-1, 3-278 TPM; Chr 2-3, 15-1324 TPM) (Table S1). The RNA-seq read alignments for both gene models show atypically high degrees of sequence disagreement with the ‘Camarosa’ reference, which suggests these transcript reads could be derived from an alternative locus such as an hypothesized deletion on Chr 2-2 (Figure S3).

In the LG 5D/Chr 5-4 QTL region, no candidate genes were found which correspond to known or hypothesized strawberry methyl anthranilate pathway components. However, a co-segregating transcript eQTL on Chr 5-4 was identified for a putative glutathione peroxidase gene (maker-Fvb5-4-augustus-gene-12.41) (transcript level *h*^*2*^=1.0) (Table 1). Single-marker analysis explains 51.1% of the transcript variation observed (AX-123358624, *p=*2.8-e9) (Figure S4A). Accumulation of this transcript positively correlates with methyl anthranilate production (Figure S4B).

A possible third methyl anthranilate signal on Chr 7 corresponds with the position of two *ANTHRANILATE SYNTHASE ALPHA* (*FanAS*-α) homoelogs (Figure S5A, Table S3). Both genes represent the only *FanAS*-α transcripts abundant in the fruit (Figure S5B), and presence/absence variation of the Chr 7-4 *FanAS*-α transcript is governed by a *cis*-eQTL which co-segregates with the methyl anthranilate signal (Figure S5C and D).

### Mesifurane QTL and Candidate Genes

Two mesifurane (*h*^*2*^=0.72) QTL were identified on LGs 1A and 7B of the FL_08-10 × 12.115-10 map (Figure 3A-B, Table 2). The mesifurane volatile QTL on LG 7B co-segregates with a transcript eQTL for the published mesifurane biosynthesis gene *O-METHYL TRANSFERASE 1* (*FanOMT1;* maker-Fvb7-4-augustus-gene-6.44*) (h*^*2*^ of transcript accumulation=0.80) (Figure 3B). Homozygosity of the mesifurane LG 7B minor allele (AX-123363184) eliminates both *FanOMT1* transcript and mesifurane production (Figure 3C). The *FanOMT1* allelic states correspond with stepwise increases in transcript abundance (Figure 3C). A novel mesifurane QTL was detected on LG 1A of the FL_08-10 × 12.115-10 genetic map (Figure 3A, Table 2). Alignments of probe nucleotide sequences to the ‘Camarosa’ genome are not subgenome-specific (Table 2).

**Table 2.** Mesifurane QTL and Candidate Gene Positions.

| Chr 1 Candidate Genes | Annotation | Fruit TPM | 'Camarosa' genome physical position |  |  |  |
| --- | --- | --- | --- | --- | --- | --- |
|  |  |  | Chr 1-1 | Chr 1-2 | Chr 1-3 | Chr 1-4 |
| augustus_masked-Fvb1-2-processed-gene-35.18 | UGT73B23/4; 96% identical | 5.3 ± 2.2 | - | 3,570,716 | - | - |
| maker-Fvb1-2-augustus-gene-137.19 | UGT85K16; 97% identical | 6.3 ± 10.3 | - | 13,720,348 | - | - |
| maker-Fvb1-4-augustus-gene-115.43 | UGT85K16 100% identical | 16.1 ± 1.3 | - | - | - | 11,555,530 |

| Chr 1 QTL | *Mesifurane*QTL *p*-value | Holiday x Korona map |  | 14.95 map |  | 'Camarosa' genome physical position |  |  |  |
| --- | --- | --- | --- | --- | --- | --- | --- | --- | --- |
|  |  | LG | Position | LG | Positon | Chr 1-1 | Chr 1-2 | Chr 1-3 | Chr 1-4 |
| AX-166520175 | 1.3E-06 | - | - | 1A | 14.522 | 26,511,819 | 4,390,575 | 1,142,702 | 1,929,385 |
| AX-166502845 | 2.4E-05 | - | - | - | - | 26,626,204 | 4,514,320 | 1,030,480 | 2,093,705 |
| AX-166520217 | 3.2E-05 | - | - | 1A | 12.229 | 27,171,481 | 4,947,347 | 560,320 | - |
| AX-166502747 | 3.2E-05 | - | - | - | - | 25,629,689 | - | 1,890,371 | 1,053,881 |
| AX-123360076 | 3.8E-05 | - | - | - | - | 25,905,230 | - | - | 1,374,796 |
| AX-166510551 | 5.2E-05 | - | - | 1A | 16.117 | - | - | - | 809,104 |
| AX-166510720 | 5.6E-05 | - | - | 1A | 14.587 | 25,788,362 | 3,789,710 | - | 1,250,466 |
| AX-89779650 | 6.1E-05 | - | - | - | - | - | 4,400,404 | - | 1,966,919 |
| AX-166510859 | 7.2E-05 | - | - | 1A | 12.381 | 27,316,667 | - | - | - |
| AX-166502836 | 1.3E-04 | - | - | - | - | - | - | - | 1,961,919 |

| Chr 7 Candidate Gene | Annotation | Fruit TPM | 'Camarosa' genome physical position |  |  |  |
| --- | --- | --- | --- | --- | --- | --- |
|  |  |  | Chr 7-1 | Chr 7-2 | Chr 7-3 | Chr 7-4 |
| maker-Fvb7-4-augustus-gene-6.44 | FanOMT1; 100% identical | 1099 ± 579 | - | - | - | 587,687 |

**Table 2 Cont.**
| Chr 7 QTL | Mesifurane<br>QTL p-value | FanOMT1<br>eQTL p-value | Holiday x Korona<br>map |  | 14.95 map |  | Chr 7-1 | Chr 7-2 | Chr 7-3 | Chr 7-4 |
| --- | --- | --- | --- | --- | --- | --- | --- | --- | --- | --- |
|  |  |  | LG | Position | LG | Positon |  |  |  |  |
| AX-166526335 | 9.4E-04 | 1.1E-05 | - | - | - | - |  |  |  | 42,976 |
| AX-166526460 | 6.6E-04 | 2.6E-05 | 7B | 60.9343 | - | - | 30,801,212 | 28,013,548 |  | 1,689,914 |
| AX-123359530 | 9.4E-04 | 1.1E-05 | 7B | 60.9343 | - | - | - | - | - | 447,154 |
| AX-123359532 | 9.4E-04 | 1.1E-05 | 7B | 60.9343 | - | - | 30,210,474 |  | 1,526,586 | 460,640 |
| AX-123359539 | 9.4E-04 | 1.1E-05 | - | - | - | - |  |  | 1,824,585 | 748,784 |
| AX-123359733 | 9.4E-04 | 1.1E-05 | - | - | - | - | - | - | - | 692,839 |
| AX-123363167 | 9.4E-04 | 1.1E-05 | 7B | 60.9343 | - | - | 32,004,806 |  | 1,170,841 | 230,562 |
| AX-123363171 | 9.4E-04 | 1.1E-05 | 7B | 60.9343 | - | - | 30,218,969 | - | 1,518,160 | 452,092 |
| AX-123363175 | 9.4E-04 | 1.1E-05 | - | - | - | - | 30,120,861 | - | 1,675,218 | 550,652 |
| AX-123366658 | 9.4E-04 | 1.1E-05 | 7B | 60.9343 | - | - | 30,190,895 | - | 1,546,336 | 475,412 |

An epistatic interaction was detected between the two QTL (F(6,173) = 13.78, *p*=9.0e-13) (Figure 3D). Homozygosity of the Chr 1-group QTL (AA genotype) strongly diminishes mesifurane abundance even when the competent Chr 7-4 *FanOMT1* allele is homozygous (BB genotype) and transcript abundance is highest (AA/BB vs. AB/BB Tukey HSD *p*=2.4e-7). At least one competent allele at each locus is required for robust mesifurane production (Figure 3D). Mesifurane abundance is somewhat higher when both alleles are homozygous (BB/BB vs AB/AB, AB/BB, and BB/AB ANOVA (F(1,149) = 4.38, *p*= 0.038).

Because the mesifurane Chr 1 QTL probe sequences align equally-well to multiple subgenomes, all Chr 1 homoeologous regions were considered for candidate gene identification. The published furaneol biosynthesis gene *FanQR* (maker-Fvb6-3-augustus-gene-21.50) is not located in Chr group 1, nor were genes of similar function found in the region. The published *F. ×ananassa* furaneol glucosyltransferase gene from Yamada et al. 2019 (UGT85K16) has two putative homologs located in the ‘Camarosa’ Chr 1 group, however they are over 10Mb from the mesifurane Chr 1 QTL (100% nucleotide identity, maker-Fvb1-4-augustus-gene-115.43; 97% nucleotide identity, maker-Fvb1-2-augustus-gene-137.19) (Table 2). The two most active *F. ×ananassa* furaneol glucosyltransferases from Song et al. 2016 (UGT71K3a/b, UGT73B23/4) have putative orthologs outside of the Chr 1 group (98% nucleotide identity, augustus_masked-Fvb3-4-processed-gene-50.19; 100% nucleotide identity; augustus_masked-Fvb2-2-processed-gene-195.2). However, UGT73B23/4 has a highly-identical second homolog in Chr 1-2 (augustus_masked-Fvb1-2-processed-gene-35.18; 96% nucleotide identity, 100% coverage) located 164 genes (0.9Mb) from the most-significant mesifurane marker (AX-166520175, *p*=2.4e-6) and 49 genes (0.2Mb) from the marker AX-166510720 (*p*=5.6e-05) (Figure 3A; Table 2).

In diverse tissues of the ‘Camarosa’ plant (AX-166520175=BB), the candidate Chr 1-2 furaneol glucosyltransferase transcript levels are high in roots but low in the ripe fruit, with fruit expression somewhat increasing with ripening series (Table S3). In the mature fruit RNA-seq populations, the Chr 1-2 candidate is modestly expressed in the fruit (5.3 ± 2.2 TPM).

Using a high-resolution melting (HRM) assay in two separate crosses (n=72), two HRM markers targeting SNPs in the mesifurane Chr 1-2 QTL (AX-166520175, AX-166502845) were tested for association with mesifurane abundance in marker-assisted seedling selection. Both Chr 1 markers were confirmed to predict the segregation of mesifurane abundance (F(1,71) = 8.25823, *p*= .0006) (Figure S6).

### Terpene QTL and Candidate Genes

A shared QTL for the production of nine terpene compounds was discovered, corresponding to Chr 3 of the *F. vesca* physical map. Only two shared terpene markers are positioned in an octoploid genetic map (AX-166504318, AX-166521725), both of which correspond to LG 3B in ‘Holiday’ × ‘Korona’ (Figure 4A, Table 3) which represents Chr 3-3 in the ‘Camarosa’ genome (Hardigan et al. 2020). For linalool (*h*^*2*^=0.749), the most-correlated QTL marker (AX-166513106, *p*= 4.2e-11) explains 14.8% of the observed variance in linalool abundance. This represents a large absolute difference as linalool is among the most abundant volatiles in strawberry fruit, commonly exceeding 100 ng^1^ gFW^-1^ hr^-1^ in the cultivars used as parental lines for these populations (Schwieterman 2013). The linalool QTL is shared with the monoterpenes 3-carene (*R*^2^=0.09, *p*=1.2e-8), β-phellandrene (*R*^2^=0.21, *p*=7.0e-7), α-limonene (*R*^2^=0.06, *p*=7.0e-7), and linalool oxide (*R*^2^=0.08, *p*=4.4e-6), and the sesquiterpenes nerolidol (*R*^2^=0.09, *p*=4.3e-6), α-caryophellene (*R*^2^=0.10, *p*=8.9e-6), α-farnesene (*R*^2^=0.16, *p*=1.6e-6), and β-farnesene (*R*^2^=0.12, *p*=2.4e-7) (Figure 4A-B, Table 4). The four sesquiterpene QTL are comprised of two shared significant markers, which are also common to the monoterpenes (Table 4). Mono- and sesquiterpene volatile abundances are very strongly correlated within-group, and moderately correlated between groups (Figure 5).

**Table 3.**
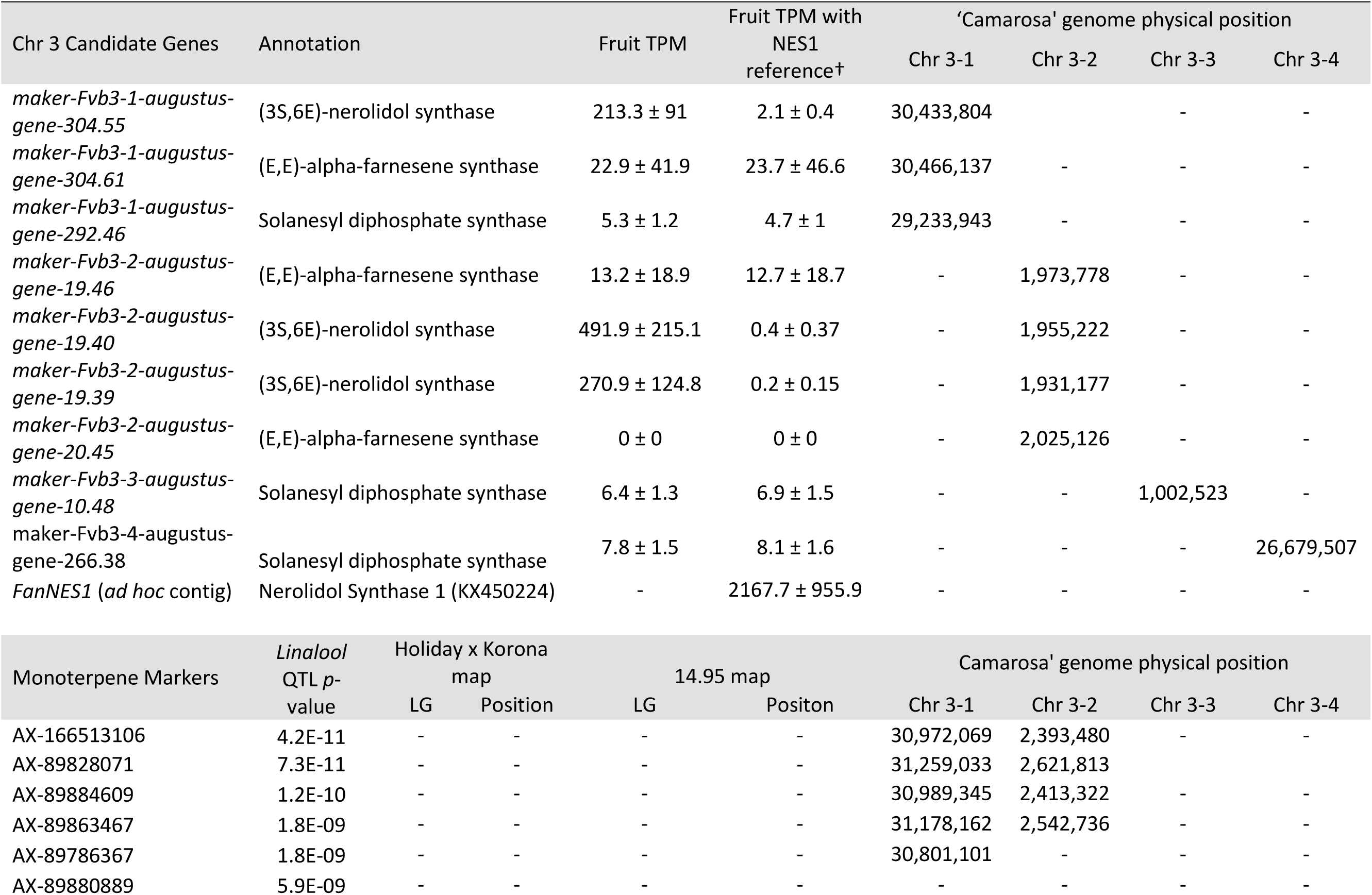
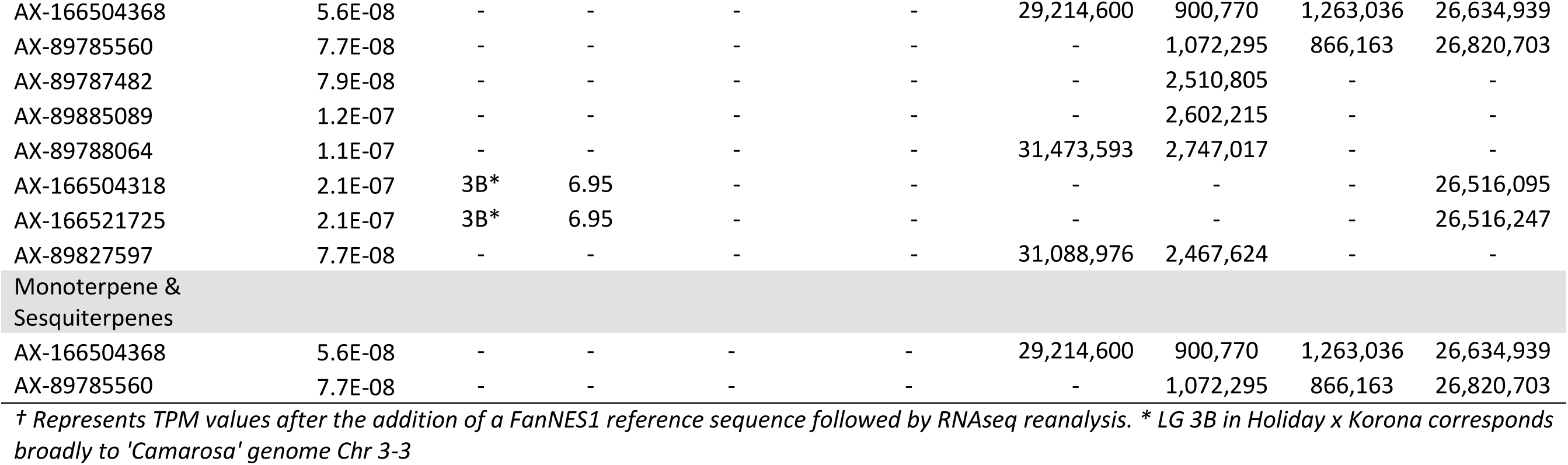
Terpene QTL and Candidate Gene Positions.

| Chr 3 Candidate Genes | Annotation | Fruit TPM | Fruit TPM with NES1 reference† | 'Camarosa' genome physical position |  |  |  |
| --- | --- | --- | --- | --- | --- | --- | --- |
|  |  |  |  | Chr 3-1 | Chr 3-2 | Chr 3-3 | Chr 3-4 |
| *maker-Fvb3-1-augustus-gene-304.55* | (3S,6E)-nerolidol synthase | 213.3 ± 91 | 2.1 ± 0.4 | 30,433,804 |  | - | - |
| *maker-Fvb3-1-augustus-gene-304.61* | (E,E)-alpha-farnesene synthase | 22.9 ± 41.9 | 23.7 ± 46.6 | 30,466,137 | - | - | - |
| *maker-Fvb3-1-augustus-gene-292.46* | Solanesyl diphosphate synthase | 5.3 ± 1.2 | 4.7 ± 1 | 29,233,943 | - | - | - |
| *maker-Fvb3-2-augustus-gene-19.46* | (E,E)-alpha-farnesene synthase | 13.2 ± 18.9 | 12.7 ± 18.7 | - | 1,973,778 | - | - |
| *maker-Fvb3-2-augustus-gene-19.40* | (3S,6E)-nerolidol synthase | 491.9 ± 215.1 | 0.4 ± 0.37 | - | 1,955,222 | - | - |
| *maker-Fvb3-2-augustus-gene-19.39* | (3S,6E)-nerolidol synthase | 270.9 ± 124.8 | 0.2 ± 0.15 | - | 1,931,177 | - | - |
| *maker-Fvb3-2-augustus-gene-20.45* | (E,E)-alpha-farnesene synthase | 0 ± 0 | 0 ± 0 | - | 2,025,126 | - | - |
| *maker-Fvb3-3-augustus-gene-10.48* | Solanesyl diphosphate synthase | 6.4 ± 1.3 | 6.9 ± 1.5 | - | - | 1,002,523 | - |
| *maker-Fvb3-4-augustus-gene-266.38* | Solanesyl diphosphate synthase | 7.8 ± 1.5 | 8.1 ± 1.6 | - | - | - | 26,679,507 |
| *FanNES1 (ad hoc contig)* | Nerolidol Synthase 1 (KX450224) | - | 2167.7 ± 955.9 | - | - | - | - |

| Monoterpene Markers | *Linalool* QTL *p*-value | Holiday x Korona map |  | 14.95 map |  | Camarosa' genome physical position |  |  |  |
| --- | --- | --- | --- | --- | --- | --- | --- | --- | --- |
|  |  | LG | Position | LG | Position | Chr 3-1 | Chr 3-2 | Chr 3-3 | Chr 3-4 |
| AX-166513106 | 4.2E-11 | - | - | - | - | 30,972,069 | 2,393,480 | - | - |
| AX-89828071 | 7.3E-11 | - | - | - | - | 31,259,033 | 2,621,813 |  |  |
| AX-89884609 | 1.2E-10 | - | - | - | - | 30,989,345 | 2,413,322 | - | - |
| AX-89863467 | 1.8E-09 | - | - | - | - | 31,178,162 | 2,542,736 | - | - |
| AX-89786367 | 1.8E-09 | - | - | - | - | 30,801,101 | - | - | - |
| AX-89880889 | 5.9E-09 | - | - | - | - | - | - | - | - |

**Table 3 Cont.**
|  |  |  |  |  |  |  |  |  |  |
| --- | --- | --- | --- | --- | --- | --- | --- | --- | --- |
| AX-166504368 | 5.6E-08 | - | - | - | - | 29,214,600 | 900,770 | 1,263,036 | 26,634,939 |
| AX-89785560 | 7.7E-08 | - | - | - | - | - | 1,072,295 | 866,163 | 26,820,703 |
| AX-89787482 | 7.9E-08 | - | - | - | - | - | 2,510,805 | - | - |
| AX-89885089 | 1.2E-07 | - | - | - | - | - | 2,602,215 | - | - |
| AX-89788064 | 1.1E-07 | - | - | - | - | 31,473,593 | 2,747,017 | - | - |
| AX-166504318 | 2.1E-07 | 3B* | 6.95 | - | - | - | - | - | 26,516,095 |
| AX-166521725 | 2.1E-07 | 3B* | 6.95 | - | - | - | - | - | 26,516,247 |
| AX-89827597 | 7.7E-08 | - | - | - | - | 31,088,976 | 2,467,624 | - | - |
| Monoterpene & Sesquiterpenes |  |  |  |  |  |  |  |  |  |
| AX-166504368 | 5.6E-08 | - | - | - | - | 29,214,600 | 900,770 | 1,263,036 | 26,634,939 |
| AX-89785560 | 7.7E-08 | - | - | - | - | - | 1,072,295 | 866,163 | 26,820,703 |

*† Represents TPM values after the addition of a FanNES1 reference sequence followed by RNAseq reanalysis. \* LG 3B in Holiday x Korona corresponds broadly to 'Camarosa' genome Chr 3-3*

**Table 4.**
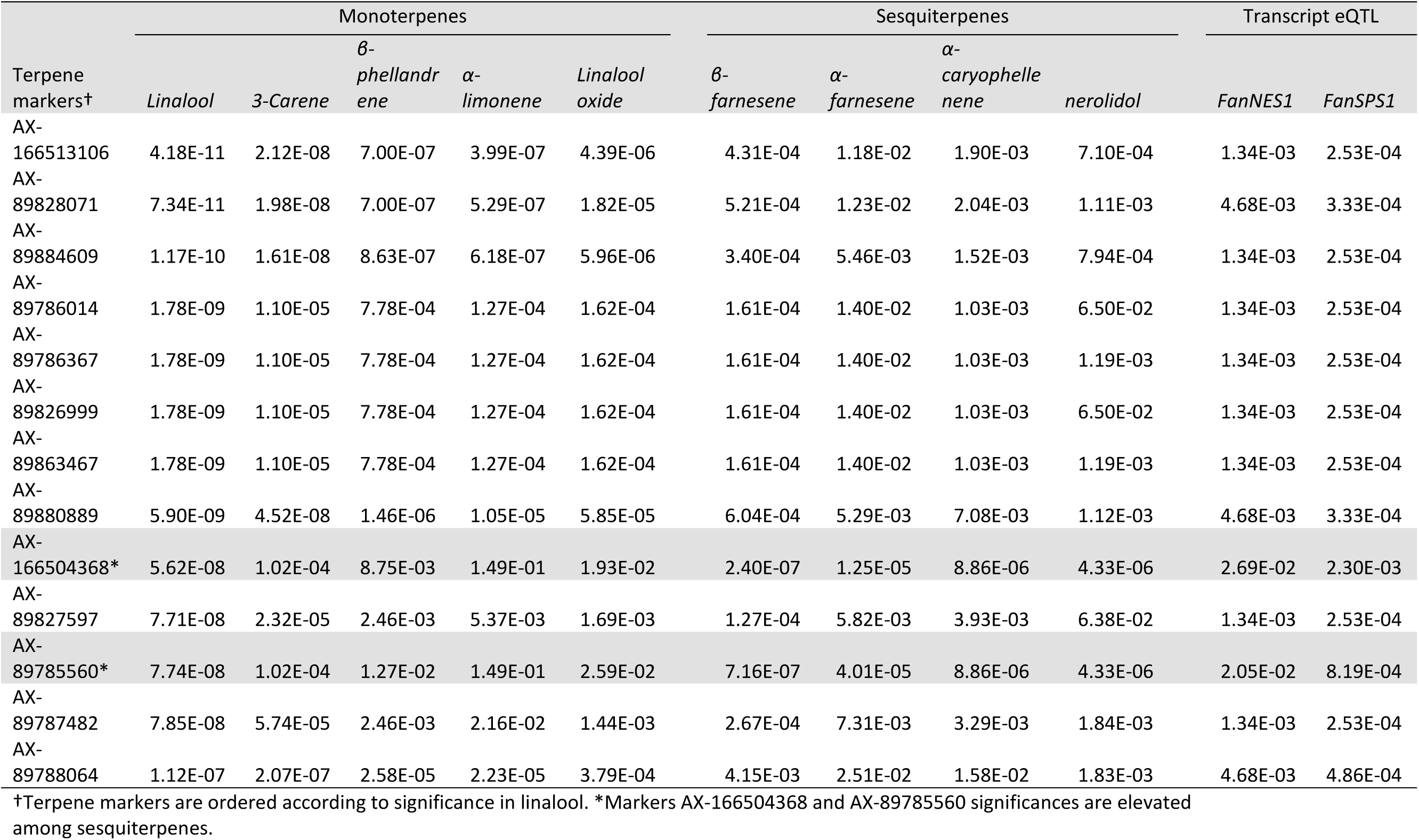
Shared Terpene QTL and Candidate Transcript eQTL Markers.

The LG 3B terpene QTL IStraw35 probe sequences align non-specifically to all four Chr 3 homoeologs (Table 3). These corresponding genomic regions contain putative terpenoid biosynthesis gene clusters, which together contain three annotated copies of *(3S,6E)-NEROLIDOL SYNTHASE*, three copies of *(E,E)-ALPHA-FARNESENE SYNTHASE*, and three copies *SOLANESYL DIPHOSPHATE SYNTHASE* (Table 3). The characterized *FanNES1* deletion responsible for linalool biosynthesis in octoploids was not detected among the ‘Camarosa’ *FanNES1*-like gene sequences, however this gene appears on Chr 3-3 in the updated ‘Camarosa’ v2 genome (Hardigan, personal communication).

To assess terpenoid-related transcript levels in fruit, the published *FanNES1* gene sequence was added *ad hoc* to the ‘Camarosa’ v1 genome as an independent contig prior to reference-based RNA-seq re-assembly. The resulting *FanNES1* transcript was the only *NES1*-like transcript abundant in fruit (Table 3). Two eQTL signals co-segregated with terpene QTL markers (Table 4). These eQTL correspond to the ambiguously-located *FanNES1* gene and to a *cis*-eQTL for a novel *SOLANESYL DIPHOSPHATE SYNTHASE* (*FanSPS*) gene on Chr 3-3 (maker-Fvb3-3-augustus-gene-10.48). Linalool abundance (n=61) is statistically correlated with *FanNES1* transcript levels (*R*^2^=0.31, *p*=0.017), but a significant relationship was not detected with *FanSPS* (*R*^2^=0.20, *p* =0.14). An additional *cis*-eQTL was detected for an *(E,E)-ALPHA-FARNESENE SYNTHASE* gene on Chr 3-2 (maker-Fvb3-2-augustus-gene-19.46) (AX-166522353; *R*^2^=0.50, *p* =1.9e-7). While this terpene biosynthesis gene is positioned closely with potential IStraw35 physical positions (Table 3), these markers do not genetically co-segregate with the terpene QTL, and transcript levels are not correlated with linalool abundance (*R*^2^=0.004, *p* =0.98).

## Discussion

Many QTL were discovered for strawberry flavor and aroma compounds known to influence the human sensory experience. These QTL are derived from eight biparental crosses phenotyped across multiple seasons under a commercial cultural system in central Florida, and are likely to be useful for making genetic gains in related germplasm. Markers correlated with these traits may be used to guide breeding decisions and identify and select for alleles mediating flavor and aroma. Potential causal genes were identified via a multi-omics approach, and provide a foundation for possible gene-editing-based approaches to improved strawberry flavor. These genetic discoveries represent new opportunities for improving flavor in commercial strawberry, and advance the basic understanding of the molecular mechanisms driving fruit flavor and aroma.

### Methyl Anthranilate

Consistent with the long-standing polygenic hypothesis for methyl anthranilate production in octoploid strawberry, multiple QTL were identified for this trait. Multi-omics analysis of QTL regions implicated several likely causal genes within distinct QTL. Because many discrete loci affect methyl anthranilate levels, and because environmental interactions transiently induce wide phenotypic swings including trait presence/absence, the interactions between loci could not be reliably measured in this sample size and diverse set of crosses. However, no loci could be identified as singularly required for production. The published *FanAAMT* gene on Chr 4, which did not emerge as a QTL in this analysis, was identified solely in the context of the biparental population ‘Florida Elyana’ × ‘Mara de Bois’ (population “10.133”), which contained only 13 analyzed progeny from one cross (Pillet et al. 2017). It is possible that these differences are due to segregating genetic factors becoming fixed or lost in subsequent populations. This hypothesis is supported by the low positive rates resulting from F1 backcrosses to ‘Mara des Bois’ (Chambers, 2013), and the fact that population “10.133” does not independently support the identified QTL regions. This QTL analysis is mostly comprised of populations using the parent ‘12.115-10’, which is a descendant of ‘Mara des Bois’ that produces more methyl anthranilate than its ancestor. It is likely this breeding line has been enriched for favorable methyl anthranilate genetics. These findings might relate more to quantitative differences in methyl anthranilate abundance, rather than the genetic presence/absence which historically defines this rare trait among strawberry cultivars.

The methyl anthranilate LG2A QTL is positively correlated with the production of two other methyl ester volatiles, namely 2-hexenoic acid, methyl ester and methyl 2-methylbutyrate. Consistent with historical segregation ratios which implicate methyl anthranilate as a polygenic trait, less methyl anthranilate variance is explained by this QTL compared with the other two methyl ester volatiles. As their precursors are not closely related, a single promiscuous methyl transferase offers a parsimonious explanation. In Pillet et al 2017, moderate methyl anthranilate levels were occasionally detected in the near-absence of the published *FanAAMT* transcript, which is suggestive of the possibility of additional methyl transferases.

While hundreds of methyl transferase genes exist in the octoploid genome, only the published *FanAAMT* has experimentally-demonstrated affinity for anthranilate. Four *FanAAMT*-like transcripts were abundantly detected in mature octoploid fruit transcriptomes. Two of these expressed *FanAAMT*-like genes correspond to the QTL on Chr 2, located within two genes (Chr 2-1) and four genes (Chr 2-3) from the most-correlated QTL markers. The expressed Chr 4-1 *AAMT*-like sequence in the ‘Camarosa’ genome is the most similar to the published Chr 4 *AAMT* sequence, whose subgenomic identity was not established (Pillet et al. 2017). This gene on Chr 4-1 might be the *FanAAMT* gene in ‘Camarosa’, particularly as RNA-seq reads from fruit transcriptomes have high sequence fidelity with this gene reference (Figure S3).

Genetic mapping suggests only a single methyl anthranilate QTL for chromosome group 2, which should be located on ‘Camarosa’ Chr 2-2. However, this QTL marker region in the Chr 2-2 physical sequence is completely absent. As ‘Camarosa’ is not capable of producing methyl anthranilate, one or more required genetic elements are expected to be missing in this reference genome. Poor RNA-seq sequence agreement with the *FanAAMT*-like Chr2-1 and Chr 2-3 homoeologs suggests the correct position of these transcript reads is not in the ‘Camarosa’ genome. It is unlikely that RNA-seq reads corresponding to the published Chr 4 *FanAAMT* transcript would map falsely to the Chr 2 candidate loci, as the published sequence is the most identical to the Chr 4-1 *FanAAMT* gene, and the RNA-seq mapping criteria excludes all non-specific reads. A comparative pan-genome analysis using a methyl anthranilate-producing individual would be highly informative, and will be undertaken in the future.

No candidate genes belonging to the hypothesized methyl anthranilate pathway are located in the Chr 5-4 region of the ‘Camarosa’ reference. However a co-segregating transcript *cis*-eQTL was detected for a putative glutathione peroxidase gene. Many but not all significant markers were shared between the trait QTL and transcript *cis*-eQTL, since methyl anthranilate levels are influenced at multiple loci while the candidate transcript is under strong single locus control. In microbes, there is precedent for heme peroxidase activity catalyzing methyl anthranilate biosynthesis (Van Haandel et al. 2000), however this reaction is unlikely to proceed via a glutathione peroxidase. It is possible that this *cis*-eQTL is simply in close linkage with the actual causal gene, which was either not correctly identified or is not present in the ‘Camarosa’ reference genome.

A possible third methyl anthranilate QTL corresponds with two Chr 7 *ANTHRANILATE SYNTHASE ALPHA* (*FanAS*-α) homoelogs. Presence/absence variation of the Chr 7-4 *FanAS*-α transcript is governed by a *cis*-eQTL which co-segregates with the putative methyl anthranilate markers at this locus. The Chr 7-2 *FanAS-α* transcript also demonstrates transcript presence/absence variation, but this is apparently due mostly to non-heritable factors that are uncorrelated with Chr 7-4 transcript level variation. Although there are few methyl anthranilate positive individuals among 61 fruit transcriptomes, none of the ten individuals with zero combined *FanAS-α* expression show methyl anthranilate production. This pathway mechanism is consistent with previous findings implicating *FanAAMT* as necessary-but-not-sufficient for methyl anthranilate production (Pillet et al. 2017). The absence of anthranilate substrate in the mature fruit would help explain the observed absence of methyl anthranilate production even when *FanAAMT* transcript levels are high. Further efforts to validate this potential QTL signal are underway.

### Mesifurane

Mesifurane (2,5-Dimethyl-4-methoxy-3(2H)-furanone, or DMMF) is derived from the methylation of furaneol (4-hydroxy-2,5-dimethyl-3(2H)-furanone, or HDF) by *FanOMT1* (Wein et al. 2002; Zorrilla-Fontanesi et al. 2012). Mesifurane variance is influenced by a loss-of-function mutation in the *FanOMT1* promoter, which eliminates transcription and mesifurane production. This model was validated by the detection of a *cis-*eQTL for the published *FanOMT1* gene, which co-segregates with the Chr 7B mesifurane trait QTL. A novel mesifurane QTL was detected on Chr 1A which is in epistasis with the Chr 7B QTL. This QTL region contains a fruit-expressed furaneol glucosyl transferase, which is 95% identical to a characterized furaneol glucosyl transferase from *F. ×ananassa*. A substrate-restricting glucosyltransferase candidate is consistent with the epistatic interaction detected with the *FanOMT1* locus. Depletion of substrate via glucosylation would limit mesifurane biosynthesis regardless of high *FanOMT1* transcript levels. Conversely, elimination of the *FanOMT1* transcript would eliminate mesifurane production regardless of substrate availability. The LG 1A mesifurane QTL was subsequently confirmed using two validation populations, providing robust support for this QTL. This two-gene model for mesifurane biosynthesis in cultivated strawberry can be exploited for genetic gain via marker-assisted selection. Moderate mesifurane levels can be maintained via dual selection for heterozygous/heterozygous allelic states, and somewhat elevated mesifurane levels can be achieved via double-homozygote selection. These findings may resolve some of the outstanding questions in mesifurane genetics posed by (Cruz-Rus et al. 2017).

### Terpenes

Homoeologous terpene gene arrays were detected for nine strawberry mono- and sesquiterpene QTL, including the desirable compound linalool. In citrus, monoterpenes and sesquiterpenes co-locate to single genomic QTL containing paralogous terpene synthases (Yu et al. 2017). We identify a similar phenomenon in cultivated strawberry. This terpene hotspot contains clusters of multiple terpenoid synthase classes, in addition to homoeologous genes on three of four subgenomes. The known biosynthesis gene *FanNES1* was associated with terpene levels via trait/transcript level correlations and trait QTL/eQTL co-segregation. Solanesyl diphosphate synthase may contribute to terpene abundances as well. The *cis*-eQTL/QTL genetic association with solanesyl diphosphate synthase on Chr 3-3 helps support the subgenomic location of *FanNES1* and the shared terpenoid QTL in the ‘Camarosa’ genome, despite only two markers being genetically mapped and probe nucleotide sequences aligning to multiple subgenomes. It is possible that the influence of other terpene-related genes in this array remain undetected due to limitations in genome completeness, marker subgenome ambiguity, and/or presence-absence variation among genomes. With additional octoploid strawberry genomes for comparison and improved subgenomic genotyping tools, complex associations in octoploid strawberry will become more robust.

## ACKNOWLEDGEMENTS

We gratefully acknowledge Aristotle Koukoulidis for genomic DNA isolation, Natalia Salinas for assistance with GC/MS sample collection and preparation, Denise Tieman for assistance with GC/MS operation, Nadia Mourad for assistance compiling eQTL results, Andrew Hanson and Harry Klee for project discussion and guidance, Alan Chambers and Jeremy Pillet for RNA isolation and RNA-seq line selection, Zhen Fan for manuscript editing and discussion, and Angelita Arredondo and Kelsey Cearley for field expertise and assistance with fruit collection.

## LITERATURE CITED

Aharoni, A., A.P. Giri, F.W.A. Verstappen, C.M. Bertea, R. Sevenier et al., 2004 Gain and loss of fruit flavor compounds produced by wild and cultivated strawberry species. The Plant Cell 16 (11):3110–3131.

Anciro, A., J. Mangandi, S. Verma, N. Peres, V.M. Whitaker et al., 2018 FaRCg1: a quantitative trait locus conferring resistance to Colletotrichum crown rot caused by Colletotrichum gloeosporioides in octoploid strawberry. Theoretical and Applied Genetics.

Arroyo, F.T., J. Moreno, P. Daza, L. Boianova, and F. Romero, 2007 Antifungal Activity of Strawberry Fruit Volatile Compounds against Colletotrichum acutatum. Journal of Agricultural and Food Chemistry 55 (14):5701–5707.

Barbey, C., M. Hogshead, A.E. Schwartz, N. Mourad, S. Verma et al., 2020 The Genetics of Differential Gene Expression Related to Fruit Traits in Strawberry (Fragaria ×ananassa). Frontiers in Genetics 10:1317.

Barbey, C., S. Lee, S. Verma, K.A. Bird, A.E. Yocca et al., 2019 Disease Resistance Genetics and Genomics in Octoploid Strawberry. G3: Genes|Genomes|Genetics:g3.400597.402019.

Bassil, N.V., T.M. Davis, H. Zhang, S. Ficklin, M. Mittmann et al., 2015 Development and preliminary evaluation of a 90 K Axiom® SNP array for the allo-octoploid cultivated strawberry Fragaria× ananassa. BMC genomics 16 (1):155.

Benjamini, Y., and Y. Hochberg, 1995 Controlling the False Discovery Rate: A Practical and Powerful Approach to Multiple Testing. Journal of the Royal Statistical Society. Series B (Methodological) 57 (1):289–300.

Bood, K.G., and I. Zabetakis, 2002 The Biosynthesis of Strawberry Flavor (II): Biosynthetic and Molecular Biology Studies. Journal of Food Science 67 (1):2–8.

Box, G.E.P., and D.R. Cox, 1964 An analysis of transformations. Journal of the Royal Statistical Society: Series B (Methodological) 26 (2):211–243.

Chambers, A., J. Pillet, A. Plotto, J. Bae, V. Whitaker et al., 2014 Identification of a Strawberry Flavor Gene Candidate Using an Integrated Genetic-Genomic-Analytical Chemistry Approach. BMC Genomics 15:217.

Chambers, A.H., 2013 Strawberry flavor: from genomics to practical applications: University of Florida.

Cruz-Rus, E., R. Sesmero, J.A. Ángel-Pérez, J.F. Sánchez-Sevilla, D. Ulrich et al., 2017 Validation of a PCR test to predict the presence of flavor volatiles mesifurane and γ-decalactone in fruits of cultivated strawberry (Fragaria × ananassa). Molecular Breeding 37 (10):131.

Edger, P.P., T.J. Poorten, R. VanBuren, M.A. Hardigan, M. Colle et al., 2019 Origin and evolution of the octoploid strawberry genome. Nature Genetics 51 (3):541–547.

Eggink, P.M., Y. Tikunov, C. Maliepaard, J.P.W. Haanstra, H. de Rooij et al., 2014 Capturing flavors from Capsicum baccatum by introgression in sweet pepper. Theoretical and Applied Genetics 127 (2):373–390.

Faedi, W., F. Mourgues, and C. Rosati, 2002 STRAWBERRY BREEDING AND VARIETIES: SITUATION AND PERSPECTIVES, pp. 51-59. International Society for Horticultural Science (ISHS), Leuven, Belgium.

Fletcher, S.W., 1917 strawberry in North America.

Folta, K.M., and H.J. Klee, 2016 Sensory sacrifices when we mass-produce mass produce. Horticulture Research 3:16032.

Hardigan, M.A., M.J. Feldmann, A. Lorant, K.A. Bird, R. Famula et al., 2020 Genome Synteny Has Been Conserved Among the Octoploid Progenitors of Cultivated Strawberry Over Millions of Years of Evolution. Frontiers in Plant Science 10:1789.

Larsen, M., and L. Poll, 1992 Odour thresholds of some important aroma compounds in strawberries. Zeitschrift für Lebensmittel-Untersuchung und Forschung 195 (2):120–123.

Lommen, A., and H.J. Kools, 2012 MetAlign 3.0: performance enhancement by efficient use of advances in computer hardware. Metabolomics 8 (4):719–726.

Noh, Y.-H., Y. Oh, J. Mangandi, S. Verma, J.D. Zurn et al., 2018 High-throughput marker assays for FaRPc2-mediated resistance to Phytophthora crown rot in octoploid strawberry. Molecular Breeding 38 (8):104.

Pillet, J., A.H. Chambers, C. Barbey, Z. Bao, A. Plotto et al., 2017 Identification of a methyltransferase catalyzing the final step of methyl anthranilate synthesis in cultivated strawberry. BMC plant biology 17 (1):147.

R. Development Core Team, 2014 R: A language and environment for statistical computing.

Raab, T., J.A. López-Ráez, D. Klein, J.L. Caballero, E. Moyano et al., 2006 FaQR, required for the biosynthesis of the strawberry flavor compound 4-hydroxy-2,5-dimethyl-3(2H)-furanone, encodes an enone oxidoreductase. The Plant cell 18 (4):1023–1037.

Racine, J.S., 2011 RStudio: A Platform-Independent IDE for R and Sweave. Journal of Applied Econometrics 27 (1):167–172.

Rambla, J.L., A. Medina, A. Fernández-del-Carmen, W. Barrantes, S. Grandillo et al., 2017 Identification, introgression, and validation of fruit volatile QTLs from a red-fruited wild tomato species. Journal of Experimental Botany 68 (3):429–442.

Schieberle, P., and T. Hofmann, 1997 Evaluation of the Character Impact Odorants in Fresh Strawberry Juice by Quantitative Measurements and Sensory Studies on Model Mixtures. Journal of Agricultural and Food Chemistry 45 (1):227–232.

Schwieterman, M.L., 2013 Metabolite analysis, environmental factors, and a transgenic approach to understanding strawberry (Fragaria× ananassa) flavor: University of Florida.

Schwieterman, M.L., T.A. Colquhoun, E.A. Jaworski, L.M. Bartoshuk, J.L. Gilbert et al., 2014 Strawberry Flavor: Diverse Chemical Compositions, a Seasonal Influence, and Effects on Sensory Perception. PLOS ONE 9 (2):e88446.

Song, C., X. Hong, S. Zhao, J. Liu, K. Schulenburg et al., 2016 Glucosylation of 4-Hydroxy-2,5-Dimethyl-3(2H)-Furanone, the Key Strawberry Flavor Compound in Strawberry Fruit. Plant Physiology 171 (1):139.

Sánchez-Sevilla, J.F., E. Cruz-Rus, V. Valpuesta, M.A. Botella, and I. Amaya, 2014 Deciphering gamma-decalactone biosynthesis in strawberry fruit using a combination of genetic mapping, RNA-Seq and eQTL analyses. BMC genomics 15 (1):218.

Sánchez-Sevilla, J.F., J.G. Vallarino, S. Osorio, A. Bombarely, D. Posé et al., 2017 Gene expression atlas of fruit ripening and transcriptome assembly from RNA-seq data in octoploid strawberry (Fragaria?×?ananassa). Scientific Reports 7 (1):13737.

Tang, Y., X. Liu, J. Wang, M. Li, Q. Wang et al., 2016 GAPIT Version 2: An Enhanced Integrated Tool for Genomic Association and Prediction. The Plant Genome 9.

Tikunov, Y.M., S. Laptenok, R.D. Hall, A. Bovy, and R.C.H. De Vos, 2012 MSClust: a tool for unsupervised mass spectra extraction of chromatography-mass spectrometry ion-wise aligned data. Metabolomics 8 (4):714–718.

Ulrich, D., E. Hoberg, A. Rapp, and S. Kecke, 1997 Analysis of strawberry flavour – discrimination of aroma types by quantification of volatile compounds. Zeitschrift für Lebensmitteluntersuchung und -Forschung A 205 (3):218–223.

Ulrich, D., and K. Olbricht, 2013 Diversity of volatile patterns in sixteen Fragaria vesca L. accessions in comparison to cultivars of Fragaria× ananassa. Journal of Applied Botany and Food Quality 86 (1).

Ulrich, D., and K. Olbricht, 2014 Diversity of metabolite patterns and sensory characters in wild and cultivated strawberries. Journal of Berry Research 4:11–17.

Ulrich, D., and K. Olbricht, 2016 A search for the ideal flavor of strawberry-Comparison of consumer acceptance and metabolite patterns in Fragaria× ananassa Duch. Journal of Applied Botany and Food Quality 89.

Urrutia, M., J.L. Rambla, K.G. Alexiou, A. Granell, and A. Monfort, 2017 Genetic analysis of the wild strawberry (Fragaria vesca) volatile composition. Plant Physiology and Biochemistry 121:99–117.

van Dijk, T., G. Pagliarani, A. Pikunova, Y. Noordijk, H. Yilmaz-Temel et al., 2014 Genomic rearrangements and signatures of breeding in the allo-octoploid strawberry as revealed through an allele dose based SSR linkage map. BMC Plant Biology 14 (1):55.

Van Haandel, M.J.H., F.C.E. Sarabèr, M.G. Boersma, C. Laane, Y. Fleming et al., 2000 Characterization of Different Commercial Soybean Peroxidase Preparations and Use of the Enzyme for N-Demethylation of Methyl N-Methylanthranilate To Produce the Food Flavor Methylanthranilate. Journal of Agricultural and Food Chemistry 48 (5):1949–1954.

Vandendriessche, T., P. Geerts, B.N. Membrebe, J. Keulemans, B.M. NicolaÏ et al., 2013 Journeys through aroma space: a novel approach towards the selection of aroma-enriched strawberry cultivars in breeding programmes. Plant Breeding 132 (2):217–223.

Verma, S., N.V. Bassil, E. van de Weg, R.J. Harrison, A. Monfort et al., 2017 Development and evaluation of the Axiom® IStraw35 384HT array for the allo-octoploid cultivated strawberry Fragaria ×ananassa, pp. 75–82. International Society for Horticultural Science (ISHS), Leuven, Belgium.

Wein, M., N. Lavid, S. Lunkenbein, E. Lewinsohn, W. Schwab et al., 2002 Isolation, cloning and expression of a multifunctional O-methyltransferase capable of forming 2,5-dimethyl-4-methoxy-3(2H)-furanone, one of the key aroma compounds in strawberry fruits. The Plant Journal 31 (6):755–765.

Whitaker, V.M., T. Hasing, C.K. Chandler, A. Plotto, and E. Baldwin, 2011 Historical trends in strawberry fruit quality revealed by a trial of University of Florida cultivars and advanced selections. HortScience 46 (4):553–557.

Yamada, A., K.i. Ishiuchi, T. Makino, H. Mizukami, and K. Terasaka, 2019 A glucosyltransferase specific for 4-hydroxy-2,5-dimethyl-3(2H)-furanone in strawberry. Bioscience, Biotechnology, and Biochemistry 83 (1):106–113.

Yu, Y., J. Bai, C. Chen, A. Plotto, Q. Yu et al., 2017 Identification of QTLs controlling aroma volatiles using a ‘Fortune’ x ‘Murcott’ (Citrus reticulata) population. BMC genomics 18 (1):646–646.

Zorrilla-Fontanesi, Y., J.-L. Rambla, A. Cabeza, J.J. Medina, J.F. Sánchez-Sevilla et al., 2012 Genetic Analysis of Strawberry Fruit Aroma and Identification of O-Methyltransferase FaOMT as the Locus Controlling Natural Variation in Mesifurane Content. Plant Physiology 159 (2):851.

