## Supplementary Figures for "Genetic Analysis of Methyl Anthranilate, Mesifurane, Linalool and Other Flavor Compounds in Cultivated Strawberry (*Fragaria* ×*ananassa*)"

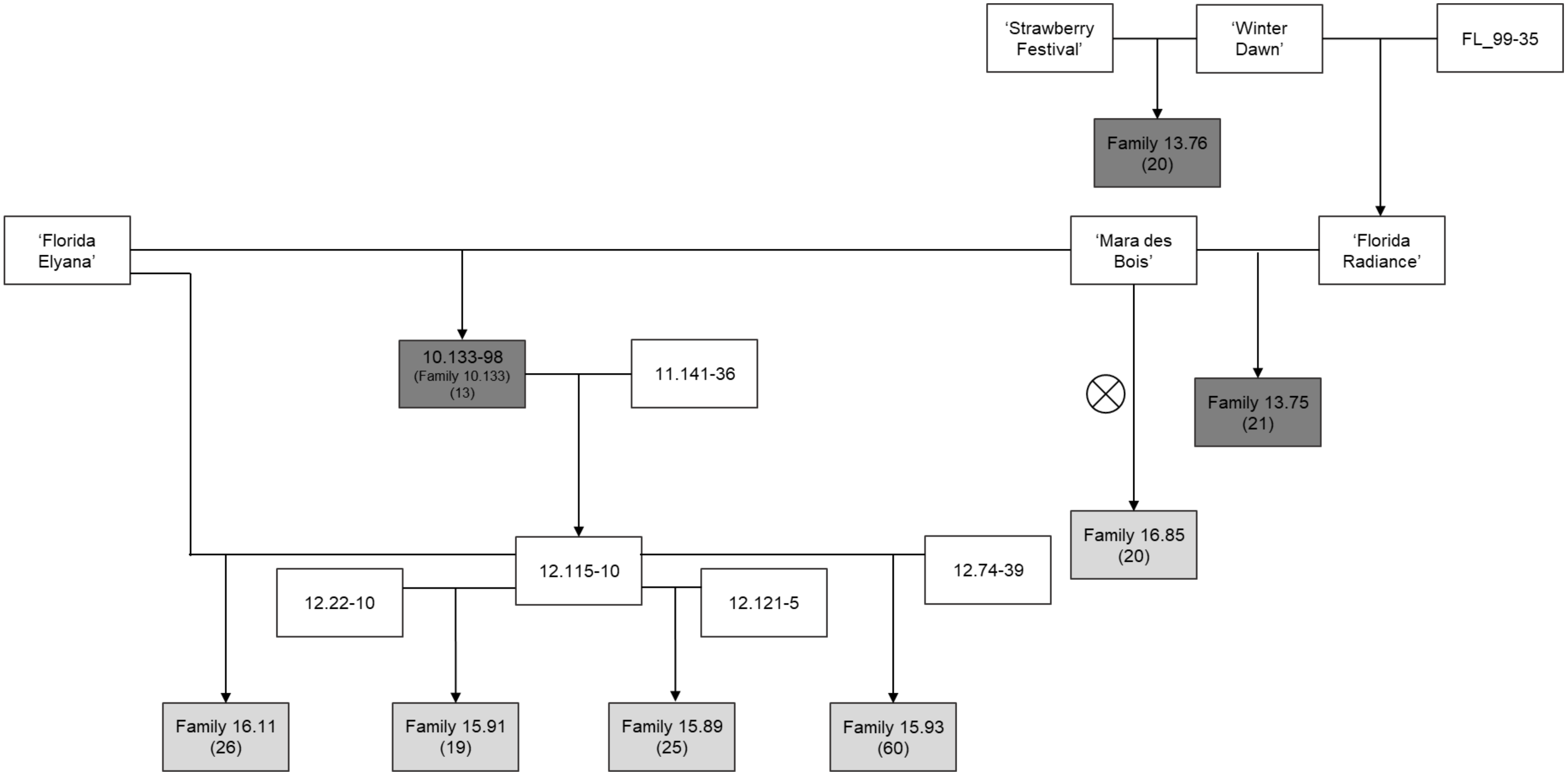


**Figure S1. Pedigree of eight interrelated strawberry families segregating for flavor and aroma.** Families used in volatile QTL analysis (light grey) are indicated with the number of analyzed progeny in parenthesis. Families used in both volatile QTL analysis and fruit RNA-seq analysis are also shown (dark gray).

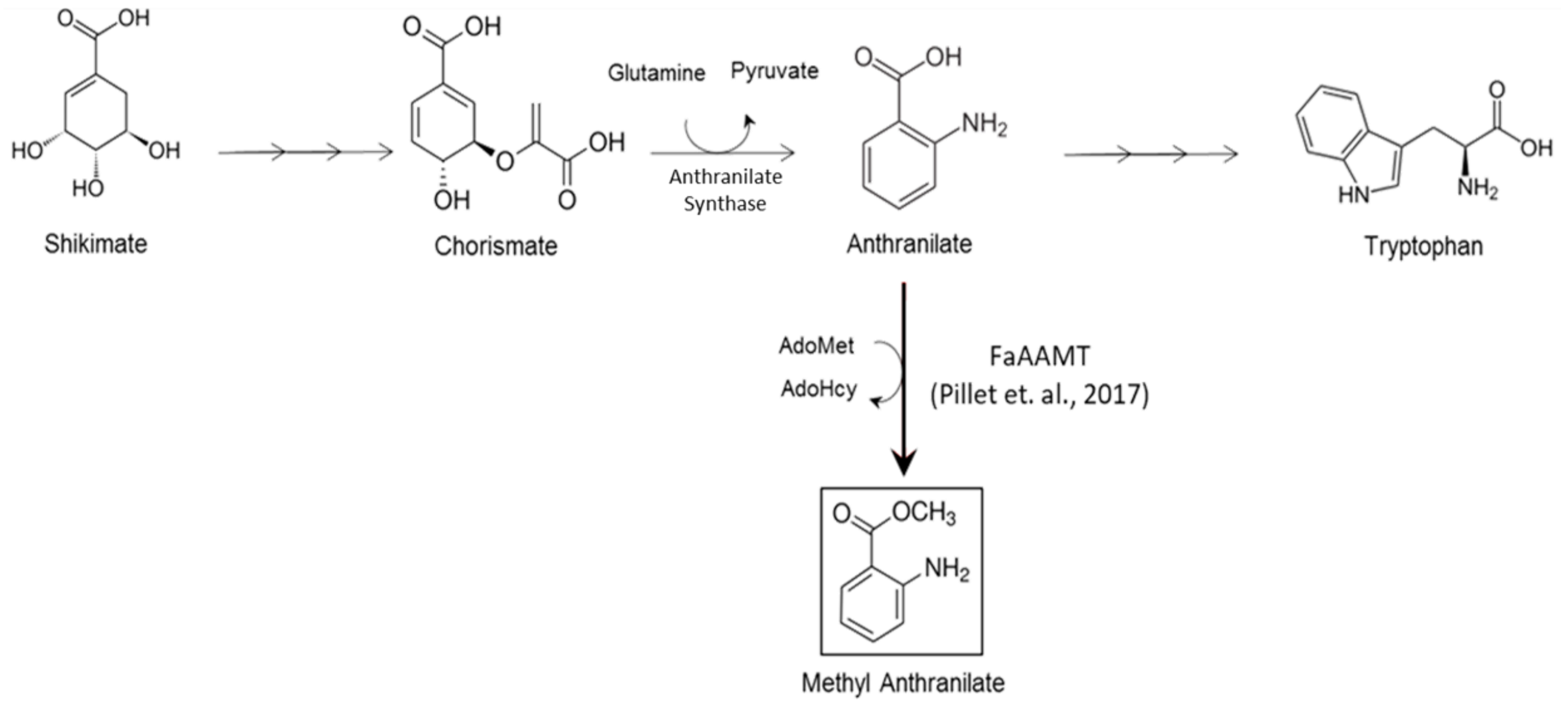


**Figure S2. The Known Methyl Anthranilate Pathway in Strawberry**. Methyl anthranilate is conditionally derived from the methylation of anthranilate (bold arrow) in the mature fruit. Anthranilate (also refered to as anthranilic acid) is derived from chorismate via the anthranilate synthase enzyme complex, and is a substrate in tryptophan biosynthesis.

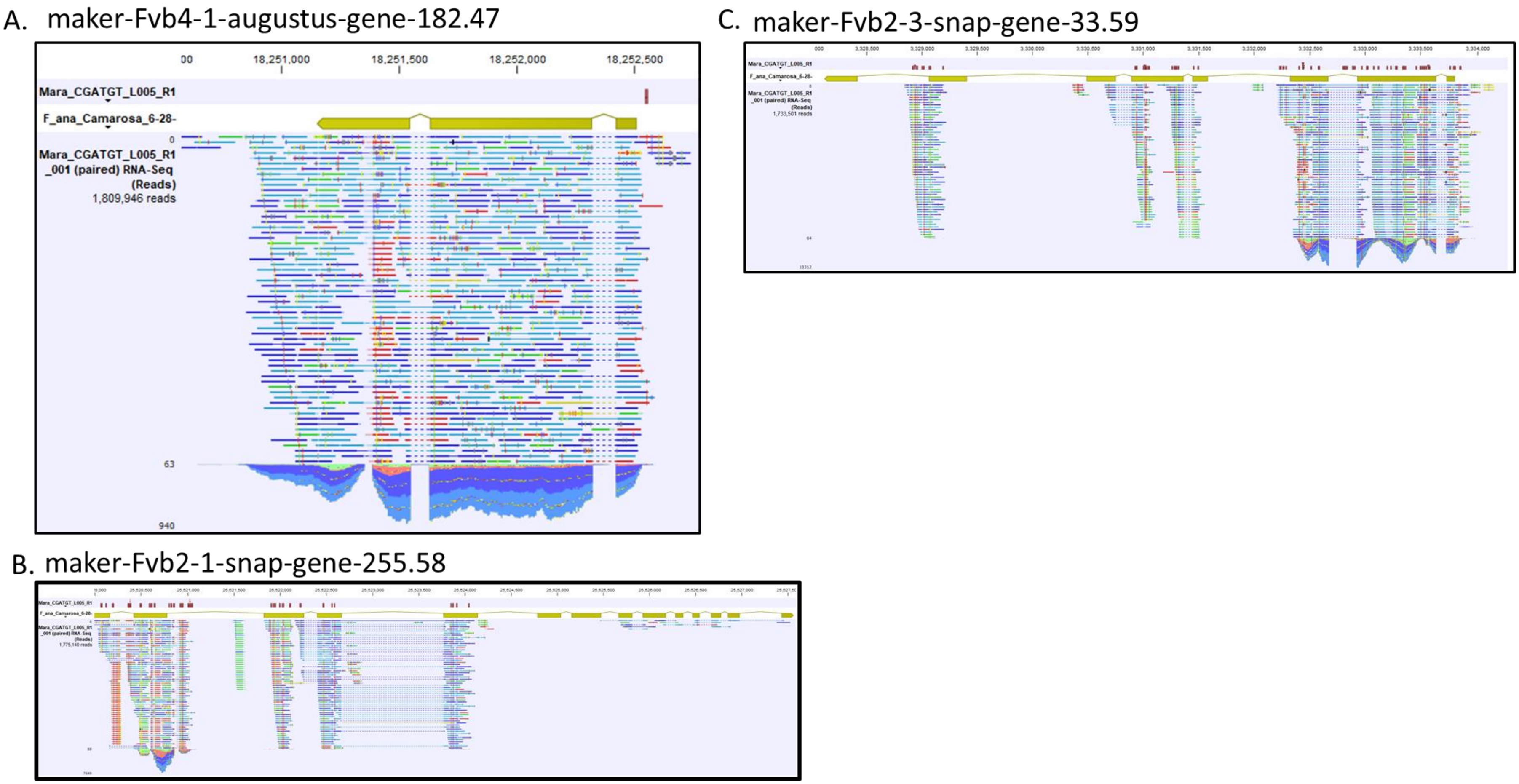


**Figure S3. RNA-seq Mapping of *FanAAMT*-like genes in 'Mara des Bois' fruit.** RNAseq read-map assemblies are shown for AAMT-like coding sequences in the 'Camarosa' octoploid genome (yellow arrows) with predicted SNP variants (red marker). **(A)** The Chr 4-1 *FanAAMT* genes shows no predicted coding sequence polymorphisms, while **(B)** the Chr 2-1 *FanAAMT* candidate gene and the **(C)** Chr 2-3 *FanAAMT* candidate gene references show poor agreement with transcript data.

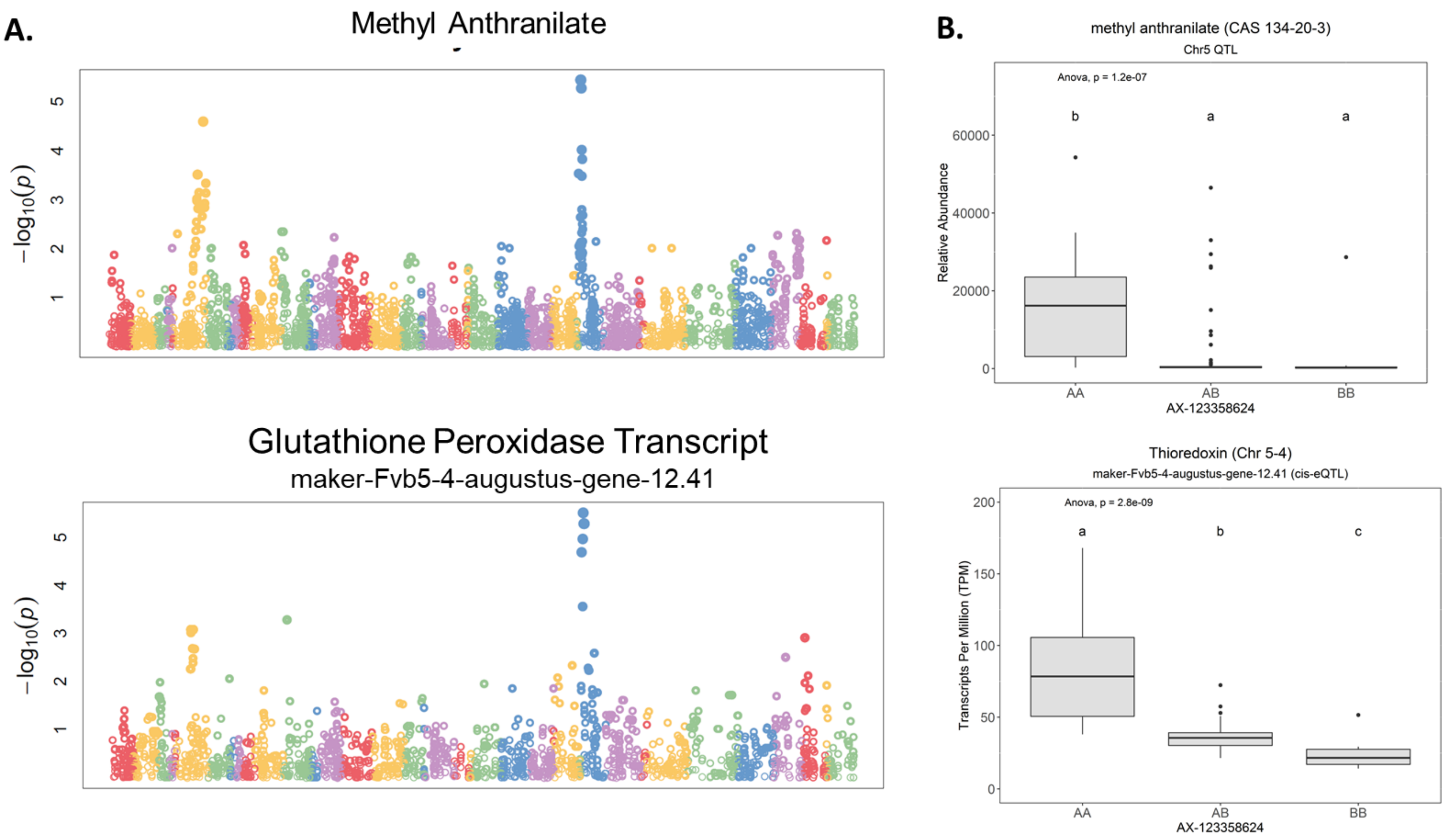


**Figure S4. Methyl Anthranilate Ch5 QTL and Candidate Genes. (A)** The methyl anthranilate QTL on chromosome 5-4 (LG 5A) is shared with a *cis*-eQTL for a putative *glutathione peroxidase* transcript. **(B)** The range of both methyl anthranilate ($r^2$ = 0.181, $p$ = 1.5e-5) abundance and *GLUTATHIONE PEROXIDASE* ($r^2$ = 0.511, $p$ = 2.8-e9) transcript abundance is shown for the shared marker AX-123358624.

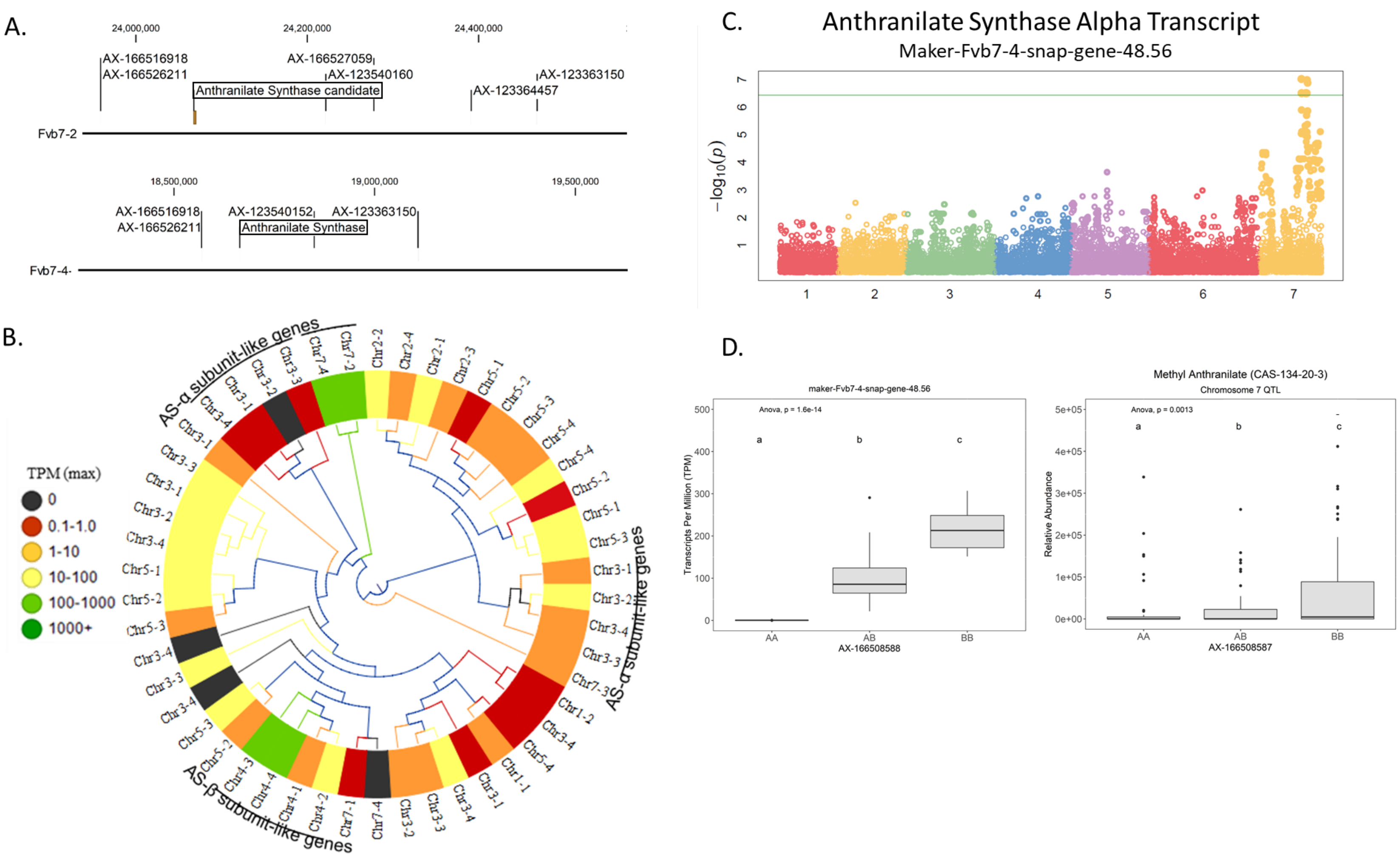


**Figure S5. Methyl Anthranilate Chr7 Candidate Genes. (A)** The putative Chr 7 Methyl Anthranilate signal corresponds to two homoeologous regions containing Anthranilate Synthase genes on Chr 7-2 (top) and Chr 7-4 (bottom). **(B)** Anthranilate Synthase-like deduced proteins in the 'Camarosa' genome are shown in a neighbor-joining cladogram, with transcript abundance heatmaps representing the highest TPM detected among the fruit transcriptomes. The Anthranilate Synthase Alpha subunit candidate genes on Chr 7-2 and Chr 7-4 are highly abundant in the fruit, as are two corresponding beta subunits genes. **(C)** Variable transcript levels of one Anthranilate Synthase candidate (Chr 7-4) are governed by a transcript *cis*-eQTL. (**D)** The eQTL for the Anthranilate Synthase alpha candidate, which governs transcript presence/absence in the fruit (left), also co-segregates with the Methyl Anthranilate Chr 7 putative QTL (right).

A.

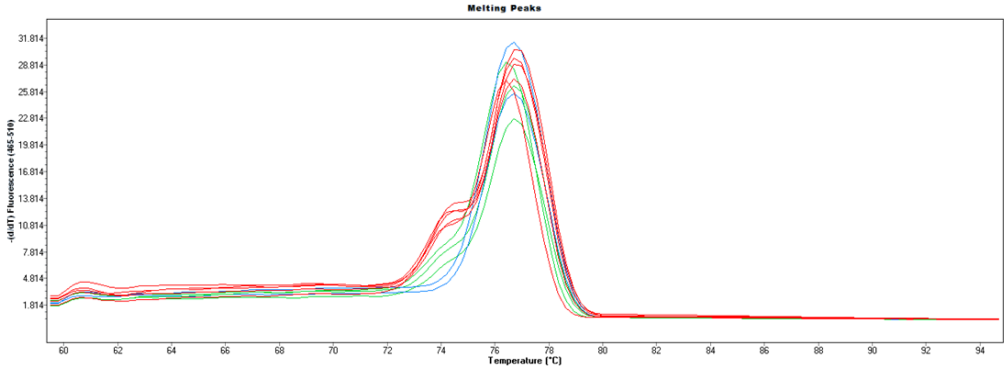


B.

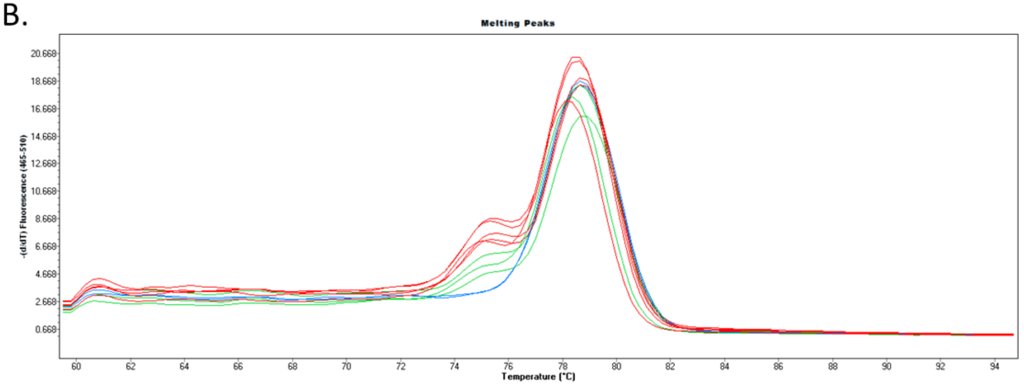


C.

| Summary of Data | | | | | | |
|---|---|---|---|---|---|---|
| | *Treatments* | | | | | |
| | 1 | 2 | 3 | 4 | 5 | Total |
| N | 16 | 40 | 16 | | | 72 |
| ∑X | 3.9201 | 6.6582 | 0.3866 | | | 10.9649 |
| Mean | 0.245 | 0.1665 | 0.0242 | | | 0.152 |
| $\sum X^2$ | 1.4135 | 2.3563 | 0.0138 | | | 3.7836 |
| Std.Dev. | 0.1738 | 0.1789 | 0.0173 | | | 0.1725 |

| Result Details | | | | |
|---|---|---|---|---|
| *Source* | *SS* | *df* | *MS* | |
| Between-treatments | 0.4082 | 2 | 0.2041 | *F* = 8.25823 |
| Within-treatments | 1.7055 | 69 | 0.0247 | |
| Total | 2.1138 | 71 | | |

**Figure S6. High-Resolution Melting (HRM) Curves for Two Chr 1 Mesifurane QTL Markers.** Ten individuals were initially confirmed to be either homozygous negative (red), heterozygous (green), or homozygous positive (blue) for the markers **(A)** AX-166520175 and **(B)** AX-166502845 based on melting curve properties. (**C)** ANOVA test statistics of fruit mesifurane abundance levels among 72 additional individuals tested by HRM confirm the Chr 1 mesifurane QTL markers.
